# Interactions between evolved pesticide resistance and pesticide exposure influence immunity against pathogens

**DOI:** 10.1101/2022.02.04.479151

**Authors:** Stephanie S.L. Birnbaum, Nora K.E. Schulz, Ann T. Tate

**Affiliations:** Department of Biological Sciences, Vanderbilt University; Institute for Evolution & Biodiversity, University of Münster

**Keywords:** experimental evolution, RNAseq, stress, pathogens, xenobiotics

## Abstract

Pesticide resistance readily evolves in natural insect populations and often coopts the same stress, detoxification, and immune pathways involved in physiological responses against primary pathogen and pesticide exposure. As a result, resistance evolution can alter antagonism or facilitation among chemical and biological pest control strategies in directions that remain difficult to predict. To investigate the interactive effects of chemical pesticide resistance, exposure, and bacterial infection on insect phenotypes, we experimentally evolved resistance to two different classes of pesticides (organophosphates and pyrethroids) in the red flour beetle, *Tribolium castaneum*. We exposed pesticide susceptible and resistant lines to pesticides, the entomopathogen and biocontrol agent *Bacillus thuringiensis* (Bt), or both. Pesticide resistance and Bt exposure were individually associated with slower development, indicating sub-lethal fitness costs of resistance and infection, respectively. After organophosphate exposure, however, beetles developed more quickly and were more likely to survive if also exposed to Bt. We used RNAseq to examine the interactive effects of pesticide resistance, pesticide exposure, and Bt exposure on gene expression. Pyrethroid-resistant insects exhibited dampened immune responses to Bt infection relative to susceptible ones. In a similar vein, simultaneous exposure to organophosphates and Bt resulted in muted stress-associated transcriptional responses compared to exposure with only one factor. Our results suggest that direct and host-mediated indirect interactions among pathogens and pesticides may buffer the cost of exposure to host fitness-associated traits within generations but exacerbate trade-offs over evolutionary time.

## Introduction

Control of insect agricultural pests and disease vectors remains a central problem worldwide as widespread use of chemical pesticides has led to the rise of pesticide resistance amongst important insect species (Bass & Jones, 2018). Insect resistance management (IRM) and integrative pest management (IPM) strategies strive to overcome problems of evolved resistance by alternating or implementing the use of both chemical pesticides and biological control measures (Sappington & Miller, 2017; Zhu et al., 2016). Entomopathogens, including bacteria, fungi, and viruses, can complement or even replace the use of chemical pesticides by increasing the mortality rate and slowing development and reproduction in pest and vector populations (Cory, 2017; Liu et al., 2017). The physiological effects of chemical and microbial control agents are not completely independent, however, as both can activate stress, detoxification, and immune responses (James & Xu, 2012). Since the evolution of pesticide resistance also often arises through these mechanisms (Minetti et al., 2020), host responses to chemical and microbial control agents could mediate facilitation or antagonism among control strategies on both proximate and evolutionary time scales (Booton et al., 2019; Hite & Cressler, 2018; Tompkins et al., 2001; Wolz et al., 2021). A better understanding of the molecular basis and phenotypic outcomes of these interactions would improve the design and predictive power of pest and vector control strategies (Bass & Jones, 2018; Sappington & Miller, 2017) while also providing fundamental insight into the integration and evolution of organismal stress and immune responses.

Pesticides target a diverse set of physiological functions in insects ranging from neurotoxic activity to the regulation of growth and development. The mechanisms of pesticide resistance show a similarly diverse set of solutions within and among pesticide classes, including target-site modifications, increased metabolic detoxification, and cuticular modifications (Boyer et al., 2012; Rösner et al., 2020). Organophosphates (OP), which inhibit acetylcholinesterase (AChE) to overexcite cholinergic synapses (Casida & Durkin, 2013), and pyrethroids (Pyr), which disrupt voltage-gated sodium channel (vgsc) function (Davies et al., 2007), are two pesticide classes widely used in agricultural systems and against disease vectors. Target-site mutations and AChE gene duplications have been described for several OP-resistant insects (Siegfried & Scharf, 2001), while Pyr resistance has been associated with mutations in the voltage-gated sodium channel (Soderlund, 2011). Target site mutations are not the only path to resistance, however. OP and Pyr pesticides are mainly detoxified through oxidation and hydrolysis, and resistance associated with differential expression of diverse canonical detoxification genes, including cytochrome P450s, esterases, and glutathione S-transferases (GSTs), has also been described for both pesticide types (Boyer et al., 2012; Meinke et al., 2021; Oakeshott et al., 2013; Pavlidi et al., 2018; Sogorb & Vilanova, 2002). Moreover, recent studies have implicated changes in cuticle and serine endopeptidase gene expression in increased resistance to penetration by OP and Pyr pesticides (David et al., 2014; Lilly et al., 2016; Tandonnet et al., 2020; Zimmer et al., 2017).

Recent evidence suggests that the mechanisms of pesticide resistance could also impact immune and physiological responses against parasites (Minetti et al., 2020). For example, esterase-mediated pesticide resistance in *Culex pipiens* is associated with changes in immune gene expression including constitutive upregulation of antimicrobial peptide (AMP) and nitric oxide synthase (NOS) genes (Vézilier et al., 2013). Moreover, resistance-associated constitutive changes in metabolic detoxification mechanisms like cytochrome P450s or GSTs can alter concentrations of damaging reactive oxygen species (ROS) that a pathogen would encounter within the host (Minetti et al., 2020; Pang et al., 2021) and influence the success of pathogen colonization, growth, and transmission (Minetti et al., 2020; Ndo et al., 2019; Wang et al., 2016). For example, changes in cap‘n’collar transcription factor expression have been shown to alter detoxification gene expression and increase ROS levels, thereby conferring pesticide resistance in several species (Kalsi & Palli, 2017; Lu et al., 2020; Misra et al., 2011) while also modifying vector competence in *Aedes aegypti* (Bottino-Rojas et al., 2018). Populations of OP and Pyr resistant mosquito strains in possession of target-site mutant *ace-1* (AChE) and *kdr* (vgsc) alleles, respectively, support a higher prevalence of *Plasmodium falciparum* parasites, but *kdr* mutations were also associated with reduced midgut oocyst burden in infected individuals (Alout et al., 2013).

Even in the absence of evolved resistance, host exposure to pesticides may impact pathogen growth directly through contact with toxins or indirectly through the induction of insect detoxification enzymes (Duke, 2018; Minetti et al., 2020). Exposure to pesticides can also have complex effects on components of the cellular, humoral, and oxidative stress responses of host immunity (James & Xu, 2012). For example, exposure to OP pesticides has been associated with increased hemocyte numbers and phenoloxidase (PO) and encapsulation activity in wax moth (*Galleria mellonella*) and Colorado potato beetle (*Leptinotarsa decemlineata*) larvae (Dubovskiy et al., 2014). However, dual exposure to OP and a pathogenic virus in silkworm larvae (*Bombyx mori*) resulted in increased mortality and differential expression of oxidative stress and AMP genes (Gu et al., 2016). Exposure to Pyr pesticides, meanwhile, has been associated with increased melanization responses and decreased replication of *Escherichia coli* bacteria (Hauser & Koella, 2020) and decreased *P. falciparum* infection prevalence and intensity in *A. gambiae* (Kristan et al., 2018). Pyr exposure is also hypothesized to impact the production of other immune responses including serine proteases, lytic enzymes including esterases and lysozymes, and reactive oxidative stress responses (James & Xu, 2012).

Clearly, both pesticide resistance and exposure independently have important effects on insect immunity against pathogens. However, it remains an open question whether resistance and exposure influence host-microbe interactions in the same direction, and whether this influence arises through the same or different mechanisms. To address this gap in knowledge, we experimentally evolved resistance to two different classes of pesticides (OP and Pyr) in the red flour beetle (*Tribolium castaneum*), an emerging model for studies on insect genomics, immunity, and resistance (Herndon et al., 2020; Rösner et al., 2020). As a stored-product pest, *T. castaneum* may be directly or indirectly exposed to the pesticides used to combat a variety of pests that co-inhabit stored grain facilities and that impose selection for resistance.

To investigate the interactive effects of pesticide resistance, exposure, and infection, we exposed our evolved lines to *Bacillus thuringiensis* (Bt), an entomopathogenic Gram-positive bacterium that has been developed into a widely used biopesticide (Raymond, 2017; Burges & Hurst, 1977; Sansinenea, 2012). While much of the research on host-Bt interactions focuses on the lethal biocontrol aspects, natural strains also occur in the environment and vary in lethality, thereby representing a selective force within natural populations (Milutinović et al., 2014; Milutinović et al., 2013). Insect resistance to Bt commonly results from changes in specific toxin-receptor interactions (Banerjee et al., 2017; Pardo-López et al., 2013), providing little expectation of cross-resistance with chemical pesticides (Siegwart et al., 2015). However, Bt-resistance has been associated with increased susceptibility to bacteria-derived pesticides (Xiao et al., 2016), and exposure to Bt has been shown to increase susceptibility to viruses and entomopathogenic nematodes (Carvalho et al., 2021; Mastore et al., 2021; Moltini-Conclois et al., 2018). The immune responses of susceptible *T. castaneum* to oral Bt infection involve significant upregulation of a suite of immune, stress, and developmental genes (Behrens et al., 2014; Ferro et al., 2019; Greenwood et al., 2017; Tate & Graham, 2017; reviewed in Pinos et al., 2021) that potentially overlap with the response to chemical pesticides and the mechanisms of pesticide resistance. However, the mode of infection may have important implications for interactions with pesticide resistance as *T. castaneum* demonstrated contrasting patterns of expression of infection-related genes dependent on the route of infection, *i.e.,* oral compared to septic infection (Behrens et al., 2014).

To explore the complex interactions between pesticide resistance and exposure to pesticides and pathogens, we first investigated the main and interactive effects of selection regime (pesticide resistant and susceptible) and pesticide exposure on host fitness-associated phenotypes after Bt infection. Having established these phenotypes, we turned to transcriptional data to identify potential mechanisms that could explain the observed phenotypes. We first asked whether evolution regime, *i.e.*, evolved pesticide resistance, alone influences constitutive gene expression and the transcriptional response to infection in the absence of pesticides. We next included pesticide exposure into the regime-by-infection interaction to investigate whether pesticides facilitate or antagonize the host response to infection, and whether differential gene expression depends on the experimental evolution regime. For both investigations, we compared the results of OP and Pyr treatments to determine whether evolved resistance or exposure to pesticides with different physiological targets exert different effects on host-pathogen interactions. Our study provides a comprehensive window into the physiological and evolutionary processes that shape interactions among two important ecological stressors.

## Methods and Materials

### T. castaneum Control and Pesticide-selected Colony Maintenance

We used six *T. castaneum* populations (Adairville, Coffee, Dorris, RR, Snavely, WF Ware) collected from stored grain facilities in the southeast USA (Jent et al., 2019). In these environments with other stored product pests, *T. castaneum* are exposed to a variety of parasites, both within and between species in the community, and may be incidentally or intentionally exposed to a range of different pesticides. Beetle colonies had been maintained under laboratory control conditions (standard diet of whole wheat flour + 5% yeast; 30°C; 70% humidity; in the dark) for at least 20 generations prior to the start of the experiment.

For each of the six ancestral *T. castaneum* populations, we initially exposed approximately 200 larvae per population to control conditions (no pesticides) or to pesticides (organophosphate [OP] or pyrethroid [Pyr]) (Suppl. Fig. 1; 18 populations total). Initial exposure doses were calculated as one tenth the manufacturer’s recommended dose (OP [Malathion 50% EC, Southern Ag]- 0.103 mg/ml; Pyr [Demon WP, 40% cypermethrin, Syngenta]- 0.0251 mg/ml). We approximated LC50 pesticide concentrations for ancestral populations using dose response curves (Birnbaum et al., 2021), and adjusted these after the second generation (LC50: 5.14 mg/ml OP and 0.188 mg/ml Pyr). To create pesticide exposure diets, we combined 0.15 g/ml standard diet with 10 ml of pesticide solutions diluted in DI water (Milutinović et al., 2013). This diet slurry was added to 1 L plastic colony containers and allowed to dry overnight at 55°C. For each pesticide-selected population, we allowed approximately 200 pupae and larvae to develop on the pesticide diets for one week before supplementing colonies with 100ml of fresh standard diet. We transferred individuals to new pesticide exposure containers every four weeks. After six generations of selection, we tested for increased pesticide resistance in F2 larvae (reared for two generations without selection to avoid parental effects) from the pesticide-regime (here, used interchangeably with evolution-regime) populations and found these populations had significantly increased survival against pesticides compared to control-regime populations (Birnbaum et al., 2021). Meanwhile, we continued selection in all populations for another two generations (to generation 8).

### Bt Spore Culture Preparation

To orally expose beetles to Bt, we created a spore culture supernatant according to previously described protocols (Milutinović et al., 2013). Briefly, we inoculated 5 ml LB liquid media with *Bacillus thuringiensis subsp. morrisoni bv. tenebrionis* (*Btt;* obtained from the *Bacillus* Genetic Stock Center, Columbus, OH) from glycerol stock and incubated overnight at 30°C. We streaked liquid *Btt* from the overnight culture onto LB agar plates and incubated the plates overnight at 30°C. The next day, 18 individual colonies were picked and each inoculated into 5 ml of Bt medium (w/V–0.75% bacto peptone (Sigma), 0.1% glucose, 0.34% KH_2_PO_4_, 0.435% K_2_HPO_4_) with the addition of 25 µL of salt solution (w/V–2.46% MgSO_4_, 0.04% MnSO_4_, 0.28% ZnSO_4_, and 0.40% FeSO_4_) and 6.25 µL of 1M CaCl_2_•2H_2_O; these cultures and a negative control (Bt medium only) were allowed to grow overnight on a bacterial shaker at 30°C, 200 rpm. The following day, 1 ml of the liquid overnight culture was inoculated into 200 ml Bt medium with the addition of 1 ml salt solution and 50 µl calcium chloride. These cultures and a negative control were incubated for seven days in darkness on a bacterial shaker at 30°C, 200 rpm; additional 1 ml salts solution and 50 µl calcium chloride were added after four days of incubation. Spore cultures were centrifuged at 4,000 rpm for 15 minutes, washed once with insect saline, and resuspended in Millipore H_2_O. We calculated the final spore concentration of 8.62*10^9^ cells/ml by counting spores using a Hausser Scientific counting chamber.

### Pesticide and Bt Spore Oral Diet Preparation

We prepared control diet by mixing 0.15 mg/ml standard diet (described above) in DI H_2_O. We prepared pesticide diets by combining approximate LC50 pesticide dilutions in H_2_O with 0.15 mg/ml standard diet (5.14 mg/ml OP and 0.188 mg/ml Pyr). *Btt* diet was prepared by mixing the previously described 8.62*10^9^ cells/ml spore culture suspension with 0.15 mg/ml standard diet. Pesticide + *Btt* diet was prepared by diluting 9.75 µl/ml OP or 0.188 mg/ml Pyr in the spore culture suspension with 0.15 mg/ml standard diet. Oral diet plates were prepared by pipetting 50 µl of the appropriate treatment (Suppl. Fig. 1; control, control + *Btt*, OP, OP + *Btt*, Pyr, Pyr + *Btt*) into 96-well cell culture plates. Plates were covered and allowed to dry overnight at 55°C.

We investigated the effects of diet disk preparation and pesticide treatments on Bt spore germination by preparing control, *Btt,* and *Btt* + pesticide diet disks as described above and counting the number of formed colonies from each treatment (*Btt*, *Btt* + OP or *Btt* + Pyr n=10; control n=6). We conducted the spore germination on two separate days. We stored dried diet disks for four or five days respectively at 30°C in the dark and then dissolved individual disks in 1 mL ultrapure water. To initiate germination and aid in the dissolving of the disks, we heated the samples to 50°C for ten minutes and then immediately plated the samples. For the bacteria culture plates, we spread 100µl of the final dilution onto LB agar, included a negative control with only water, incubated plates over night at 30°C and counted all *Btt* colonies after 15 hours. On the first day, we cultured bacteria from a 1:10^7^ dilution in ultrapure water. On the second day, we used a 1:10^6^ dilution, to ensure that sufficient colonies would grow. We analyzed the effect of treatment on the number of colony forming units (CFUs) using a generalized linear mixed model with day as a random factor and a negative binomial distribution.

### Btt in vitro Growth Curve Assays

To measure *in vitro* effects of pesticides on Bt growth, we performed growth curve analyses for OP and Pyr exposure separately. For each analysis, *Btt* from a glycerol stock was freshly streaked onto Nutrient Broth (NB) agar plates and allowed to grow overnight at 30°C. We grew overnight cultures from single colony inoculates (n = 9) in 5 ml NB liquid media at 30°C, 200 rpm; cultures were incubated for only 16 hours to avoid entrance into sporulation phase. The following day, we grew log phase bacterial cultures by inoculating 100 µl of the overnight culture into 2.9 ml NB liquid media and incubated for 3.5 hours at 30°C, 200 rpm.

To replicate *in vivo* conditions, each growth curve analysis included control NB, 1:10 pesticide NB dilution, and 1:100 NB pesticide dilution treatments based on larval exposure OP and Pyr pesticide concentrations (described above). We added 200 µl of the appropriate treatment solution to a 96-well CYTO-One spectrophotometer plate. An aliquot of log phase bacterial culture (n = 9) was added to each well to achieve an initial OD_600_ = 0.4 (3 technical replicates/ culture). OD_600_ bacterial growth was measured every 15 minutes for 17 hours using a BioTek Epoch 2 spectrophotometer (30°C). Three or four independent growth curve replicates were performed for each pesticide.

We obtained growth curve metrics using the R package growthcurver (Sprouffske & Wagner, 2016). For OP and Pyr separately, we analyzed the individual effects of replicate and treatment on growth rate values (r) obtained from growthcurver using non-parametric Kruskal-Wallis tests. We measured the effect of pesticide treatment separately for each replicate if the effect of replicate was significant.

### Exposure of Control and Pesticide-selected Populations to Pesticide and Bt Oral Diets

After eight generations of selection, we generated same-age F2 larvae from pesticide-selected and susceptible colonies. Approximately 200 adults from Gen. 8 pesticide-selected and susceptible control colonies were allowed to reproduce on control diet for three days. We removed these F0 adults and allowed offspring to develop to F1 adults. To generate F2 experimental larvae, we placed 4-5 replicates of 80 adults per selection line in 100 mm petri dishes with control diet and allowed them to reproduce for 24 hours. F2 larvae were allowed to develop for 12 days.

We placed twelve day old larvae (∼4 mm long) on oral diet plates prepared as described in the previous section (Suppl. Fig. 1; n = 96/ population/ treatment). Half of the plate (n = 48) was monitored daily for three weeks for survival and development. Individuals in the remaining half were sampled after two days of exposure prior to the onset of mortality and used for RNAseq analyses (described below).

### Statistical Methods for Fitness Experiments

The effects of selection regime (Regime), pesticide exposure (PTx), Bt exposure (BtTx), and interactions between the three factors on beetle survival and development were investigated using population as a random factor in Cox mixed effects survival models with the package coxme (Therneau, 2015) in R. For both time to death and time to develop into adulthood, we evaluated the individual effects of pesticide exposure (PTx), selection regime (Regime), and Bt exposure (BtTx), and the interactions between selection regime and pesticide exposure (Reg:PTx), selection regime and Bt exposure (Reg:BtTx), and co-exposure to pesticides and Bt (PTx:BtTx). The full model for both time to death and adult development time is as follows: Surv(Day, Status) ∼ (PTx + BtTx + Regime + Regime:PTx + Regime:BtTx + PTx:BtTx + (1|Pop)). Hazard ratios were calculated from coxme survival models and were plotted using the sjPlot package in R (Lüdecke, 2021).

### Septic Infection Experiment

Given the impact of infection route on immune gene expression (Behrens et al., 2014), we wanted to clarify whether the interaction between pesticides and infection outcomes might be infection route-dependent. We selected two populations, Coffee and Dorris, as representative populations that may demonstrate interactions with between pesticide resistance and exposure on infection outcomes. Therefore, we exposed 14-day old larvae from control- and OP-regimes to control and OP oral diets (as described above) for three days, and then septically infected individuals with *Btt* and recorded their survival.

We produced a mixture of *Btt* cultures of vegetative cells from both the logistic growth and early stationary phase for septic infections as previously described (Jent et al., 2019), which caused around 50% mortality in preliminary experiments. We pricked larvae laterally between the head and second segment with an ultra-fine borosilicate needle dipped in the bacterial suspension and recorded survival after 24 hours. We repeated the experiment three times with 20 individuals per population (Coffee or Dorris), selection regime (OP selected and control) and pesticide treatment (OP exposure and control) each (n = 480).

Differences in survival between the control group (control selection regime, no pesticide) and the other groups were analyzed in a GLMM with experiment (block) and population as random factors and a binomial error distribution (Bates et al., 2015).

### RNA Extractions and Sequencing

Samples for 3’ TagSeq (Lohman et al., 2016) were collected after two days of exposure to oral diets for three representative populations (Coffee, Dorris, Snavely), prior to the main onset of mortality. For each population, regime, and treatment, we collected nine surviving larvae and immediately stored them at -80°C until processing, and subsequently pooled larvae into three replicates of three larvae each. We extracted total RNA using a Qiagen RNeasy Mini kit according to manufacturer protocols and confirmed nucleic acid concentration and purity using a Nanodrop. We depleted remaining DNA using the Invitrogen RNaqueous Micro DNase treatment according to manufacturer protocols. We confirmed the concentration and quality of these DNA-depleted RNA samples using a Bioanlayzer (RIN > 9). We generated single indexed 3’ TagSeq libraries from 1 μg total RNA using the Lexogen QuantSeq 3’ mRNA-Seq Library Prep Kit according to manufacturer’s protocols; libraries were amplified using 15 PCR cycles. Library concentration and quality were confirmed prior to PE100 sequencing on four runs on the NextSeq platform (VANTAGE, Vanderbilt University).

### RNAseq Analyses

Adapters were trimmed from the raw mRNAseq reads and low-quality reads were removed using fastqc (Andrews 2010); an average of 10,869,280 reads per sample were retained (Suppl. Table 1). Sequence reads are deposited in the NCBI SRA under the BioProject Accessions: PRJNA753142, PRJNA756317. We took advantage of the well-annotated, published genome for *T. castaneum* (Herndon et al., 2020) to align trimmed reads to the most recent *T. castaneum* assembly (Tcas5.2) using BWA-MEM (avg. reads mapped = 84.89%, s.d. = 0.023; Suppl. Table 1) (Li, 2013). Gene count files were generated using samtools (Li et al., 2009). We used DESeq2 (Love et al., 2014) to perform differential expression analyses.

First, to investigate the effects of Bt exposure and pesticide selection regime on gene expression in the absence of pesticide exposure, we subset only pesticide-free and pesticide-free + Bt libraries and examined the effects of selection regime, Bt exposure, and their interaction on gene expression (no pesticide model form: count matrix ∼ Reg + BtTx + Reg:BtTx). Next, we analyzed the main effects of pesticide exposure, pesticide selection regime, and Bt exposure and the interactive effects between selection regime and pesticide exposure, selection regime and Bt exposure, and pesticide and Bt exposure for OP and Pyr separately (OP and Pyr model form: count matrix ∼ PTx + Reg + BtTx + Reg:PTx + Reg:BtTx + PTx:BtTx). For each factor, the control selection regime, control pesticide treatment, or control Bt treatment were set as the baseline comparisons.

Differentially expressed transcripts were annotated using the Ensembl *T. castaneum* database and the R package ‘biomaRt’ (Durinck et al., 2009). We manually curated the annotated gene lists to categorize differentially expressed transcripts into groups potentially impacted by pesticide resistance and exposure and Bt treatment such as those involved in detoxification, immunity, development, and cuticle associated transcripts (Suppl. Table 2). Gene ontology (GO) enrichment analyses for the subsets of significantly differentially expressed gene sets were performed using the online Gene Ontology Resource (Ashburner et al., 2000; Gene Ontology Consortium, 2021).

To analyze global patterns of gene expression, we used a weighted gene co-expression network analysis with the R package ‘wgcna’ (Langfelder & Horvath, 2008). This approach allows us to group genes with correlated expression patterns into modules and statistically associate gene module expression with experimental factors. We constructed the initial network using all replicate samples for all treatments (n = 126) with a variance stabilizing transformation. A soft thresholding power of 6 was used to construct the topological overlap matrix, and we then identified modules with highly correlated module expression patterns (p < 0.05).

## Results

To investigate how evolved pesticide resistance and pesticide exposure impact host interactions with pathogens, we exposed control-regime and pesticide-selected populations to control and pesticide oral treatments, with and without Bt. We measured the main and interactive effects of pesticide selection regime (Reg), pesticide treatment (PTx), and Bt treatment (BtTx) on survival, development, and gene expression for OP and Pyr separately. In brief, we found that pesticide selection resulted in increased resistance to pesticide exposure. Exposure to *Btt* did not have an overall effect on survival but did delay development, and dual exposure to OP and *Btt* resulted in improved survival compared to larvae exposed to only OP. Moreover, dual exposure to either OP or Pyr and *Btt* resulted in faster development compared to individuals exposed to only pesticides. Last, we identified antagonistic gene expression interactions between *Btt* and OP exposure and *Btt* and Pyr selection that indicate changes in immune function in infected individuals either exposed or selected with different pesticides.

### Effects of Pesticide Selection and Exposure to Pesticides and Bt on Survival and Development

Overall, exposure to both pesticide types, OP (OP PTx) and Pyr (Pyr PTx), decreased survival likelihood, while there was no main effect of Bt treatment (BtTx) on survival (Figs. 1A-B; Suppl. Table 3). Compared to control-regime populations, pesticide selection significantly improved survival after pesticide exposure, reinforcing previous observations of evolved pesticide resistance (Figs. 1A-B, compare OP Reg:PTx and Pyr Reg:PTx; Suppl. Table 3; Birnbaum et al., 2021). Interestingly, there was a positive significant interaction between OP and Bt treatment (OP PTx:BtTx) indicating that larvae dually exposed to OP and *Btt* had improved survival compared to larvae exposed to OP only (Fig. 1A; Suppl. Table 3). There was no significant interaction between Pyr exposure and Bt treatment (Pyr PTx:BtTx) or between either pesticide selection regime and Bt treatment (OP Reg:BtTx, Pyr Reg:BtTx) on survival (Figs. 1A-B; Suppl. Table 3). Thus, while pesticide selection resulted in increased pesticide resistance, pesticide selection regime did not interact with *Btt* exposure to influence survival compared to control-regime populations. However, dual exposure to OP and *Btt* improved survival outcomes compared to pesticide treatment alone.

**Figure 1.**
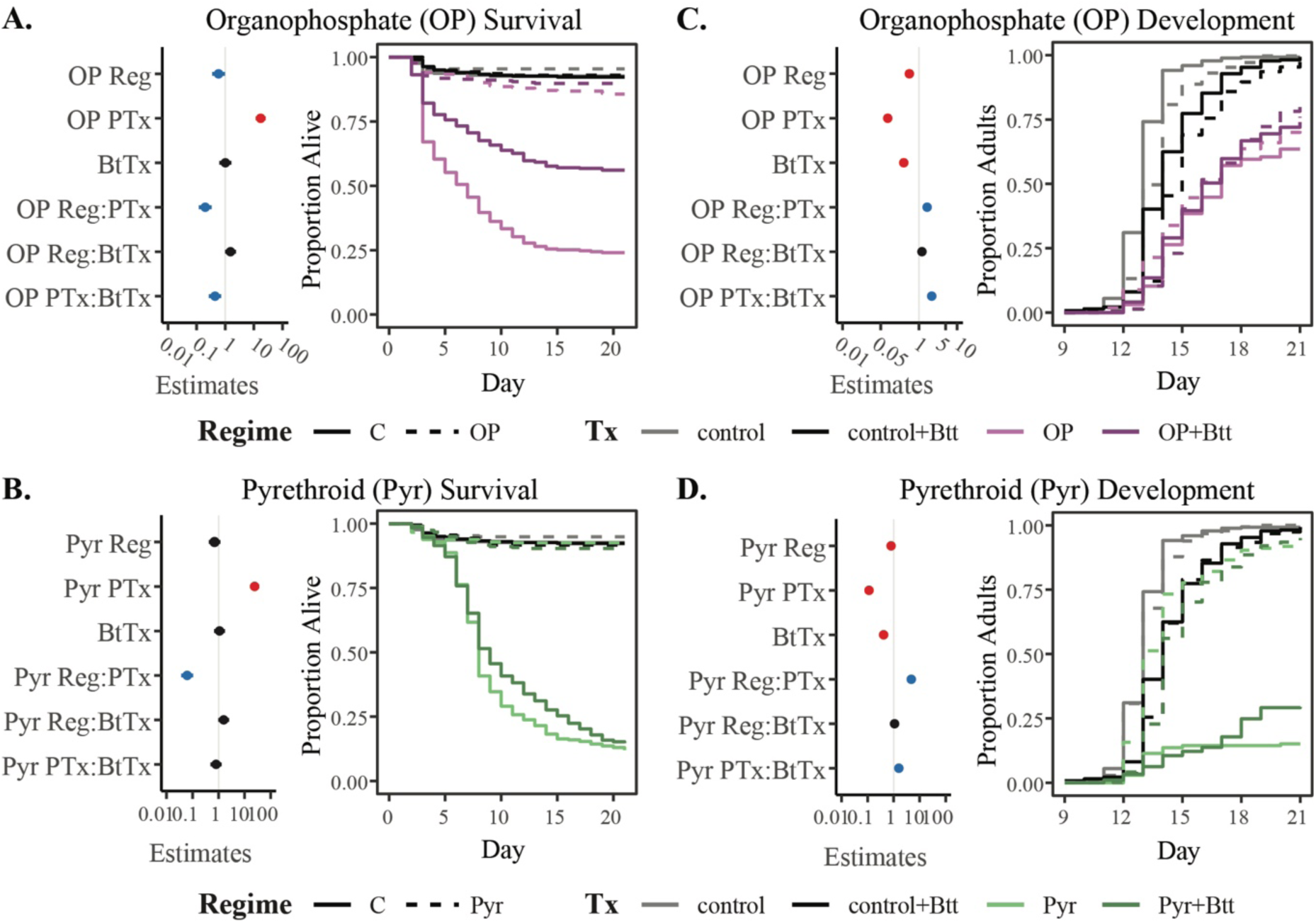
Overall effects of organophosphate (OP) and pyrethroid (Pyr) factors on survival and development. Hazard ratios depicting the effects of selection regime (OP Reg or Pyr Reg), pesticide exposure (OP PTx or Pyr PTx), Bt treatment (BtTx), the interaction between selection regime and pesticide exposure (OP Reg:PTx or Pyr Reg:PTx), the interaction between selection regime and Bt treatment (OP Reg:BtTx or Pyr Reg:BtTx), and the interaction between pesticide exposure and Bt treatment (OP PTx:BtTx or Pyr PTx:BtTx) on survival (A, B) and development (C, D) are shown in the left panels. Raw data illustrating the proportion of surviving individuals (A, B) or the proportion of adults (C, D) are shown in the right panels. For hazard ratio plots depicting survival and development estimates, positive significant effects (higher survival or faster development) are colored blue, negative significant effects (lower survival or slower development) are red, and nonsignificant effects are black (Suppl. Table 2).

Pesticide selection regime (OP Reg, Pyr Reg), pesticide exposure (OP PTx, Pyr PTx), and Bt treatment (BtTx) individually delayed development, indicating a cost from each (Figs. 1C-D; Suppl. Table 3). There was no significant interaction between pesticide regime and Bt treatment on development (OP Reg:BtTx, Pyr Reg:BtTx; Suppl. Table 3). However, resistant individuals developed faster than susceptible individuals in the presence of pesticides, as indicated by a significant interaction between pesticide regime and treatment (OP Reg:PTx, Pyr Reg:PTx; Fig. 1C-D; Suppl Table 3). There was also a significant positive interaction effect between both pesticide treatments and *Btt* exposure (OP PTx:BtTx, Pyr PTx:BtTx), indicating that larvae exposed to both *Btt* and pesticide treatments developed faster than those exposed to only pesticide or only *Btt* (Figs. 1C-D; Suppl. Table 3). Overall, exposure to pesticides delayed development but this effect was mitigated in pesticide-selected individuals. *Btt* exposure lengthened development time regardless of evolution regime but concurrent pesticide exposure partially mitigated this effect.

### Direct Effects of Pesticides on Bt

To investigate direct interactions between pesticides and Bt, we used *in vitro* growth curves and viable spore counts of oral diet disks. *In vitro Btt* growth rate significantly increased upon OP exposure, but there was no effect of Pyr exposure on *in vitro* growth rate (Suppl. Fig. 2; Suppl. Table 4). The germination rates of *Btt* spores on the diet disks exhibited no significant differences among control, OP, or Pyr pesticide diet treatments (Suppl. Fig. 3; Suppl. Table 5).

### The influence of pesticide exposure on route-specific infection outcomes

While oral infection with *Btt* did not result in immediate excess mortality compared to unexposed individuals, we considered the possibility that infection outcomes could be exposure route specific. To determine whether the interaction between pesticide resistance, exposure to pesticides, and infection outcomes might be route-dependent, we septically infected susceptible and OP-selection regime larvae exposed to control and OP diets. Individuals from the susceptible regime that were exposed to OP had lower survival against septic *Btt* infection compared to the other groups (Fig. 2; Suppl. Table 6) indicating an outsized cost of combined OP exposure and septic *Btt* infection in control regime larvae. This result contrasts with the improved survival observed in the Bt x OP treatment interaction term after oral exposure in that after septic infection, beetles co-exposed to OP do not have higher survival (Fig. 1A, OP PTx:BtTx).

**Figure 2.**
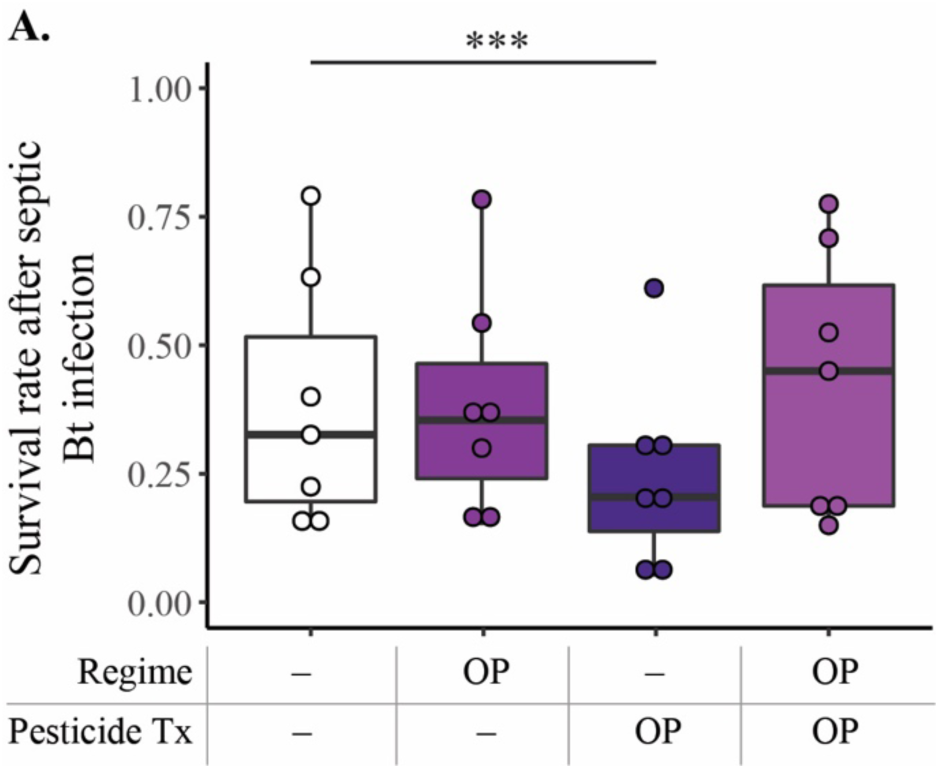
The influence of pesticide exposure on septic infection outcomes: control-regime larvae exposed to OP are more susceptible to *Btt* septic infection. The proportion of surviving individuals 24h post septic *Btt* infection are shown for control (Regime –) and OP selection regime (Regime OP) larvae exposed to control (Pesticide Tx –) or OP diet (Pesticide Tx OP) treatments. Circles indicate surviving proportions of individuals per block. Significant differences between groups are denoted with asterisks (p < 0.0005; Suppl. Table 6).

### Overall Effects of Btt Exposure, Pesticide Selection, and Pesticide Exposure on Gene Expression

Overall, we identified differentially expressed genes important in immunity in beetles exposed to *Btt* compared to unexposed beetles. We also identified differentially expressed genes implicated in detoxification and resistance processes in OP- and Pyr-regime populations compared to control-regime populations. Few genes were significantly differentially expressed in pesticide-regime populations exposed to *Btt* compared to control-regime populations. However, Pyr-regime larvae infected with *Btt* exhibited dampened expression of immune genes. Dual exposure to OP pesticides and *Btt* also resulted in dampened immune, cuticle, and detoxification gene expression compared to expression with *Btt* infection alone. We expand on these main results below.

### Effects of Btt Exposure and Pesticide Selection on Gene Expression in the Absence of Pesticide Exposure

We investigated the interaction between pesticide selection regime and *Btt* exposure on gene expression in the absence of pesticides, as this provides information on the main host response to oral *Btt* infection as well as whether the evolution of pesticide resistance modifies the host response to *Btt*. Using only the data from individuals never exposed to pesticides, we identified 104 genes that were differentially expressed in *Btt*-exposed individuals compared to unexposed individuals (Fig. 3A; Bt Tx +). This gene set largely concurred with previously published results on Bt infection in *T. castaneum* (Behrens et al., 2014; Greenwood et al., 2017; Tate & Graham, 2017), and included the upregulation of AMPs (two attacins and two defensins), a pathogenesis-related protein, and a cytochrome P450 (Fig. 4, no pesticide model BtTx). GO enrichment analyses revealed the importance of immune system processes and defense against bacteria, among other immune related terms, in the genes differentially expressed with *Btt* treatment in the absence of pesticide exposure (Suppl. Fig. 4A, no pesticide model BtTx).

**Figure 3.**
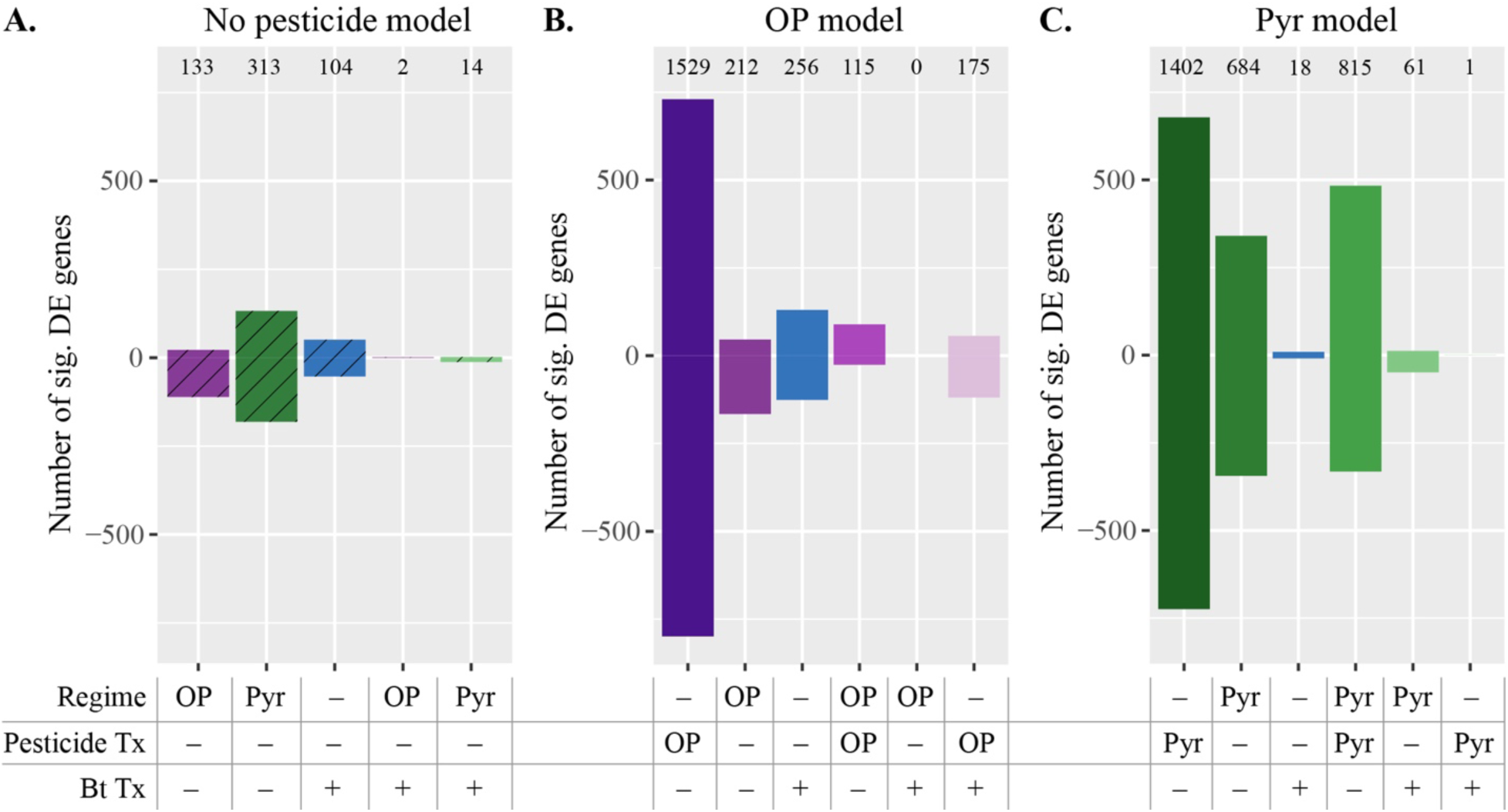
Number of significantly differentially expressed (sig. DE) genes (padj < 0.05) for factors within each DESeq2 model (number of sig. DE transcripts are denoted above bars; relative to control regime or no pesticide/ Bt treatments, positive counts are upregulated transcripts and negative counts are downregulated). A. No pesticide model: main and interaction effects of pesticide selection regime and Bt treatment in the absence of pesticide exposure. Hatched bars indicate no pesticide treatment in the model. B. OP model: main and interaction effects of OP selection regime, pesticide exposure, and Bt treatment. C. Pyr model: main and interaction effects of Pyr selection regime, pesticide exposure, and Bt treatment. The table below indicates the differentially expressed factor. Main effect of selection regimes = Regime OP/ Pyr. Main effect of Bt treatment = Bt Tx +. Main effect of pesticide treatments = Pesticide Tx OP/ Pyr. Interactions between selection regimes and pesticide exposure = Regime OP/ Pyr & Pesticide Tx OP/ Pyr. Interactions between selection regimes and Bt exposure = Regime OP/ Pyr & Bt Tx +. Interactions between pesticide and Bt exposure = Pesticide Tx OP/ Pyr & Bt Tx +.

**Figure 4.**
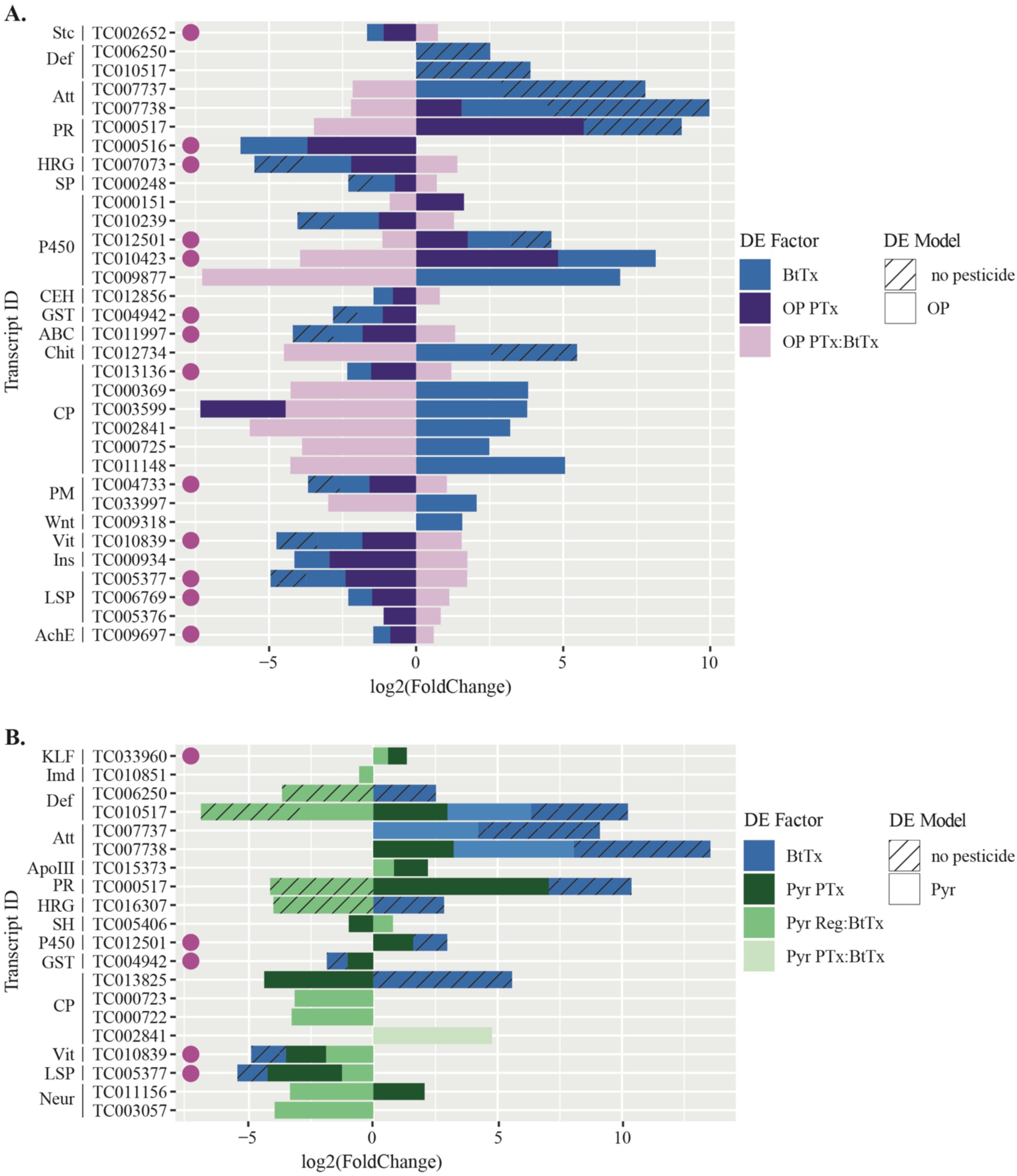
Profiles of differentially expressed transcripts demonstrate opposing responses in Bt-infected individuals that were or were not also exposed to or selected with pesticides. Transcripts significantly differentially expressed (padj < 0.05) with Bt treatment and interactions with **A.** OP exposure, and **B.** Pyr selection regime and exposure. The DE Factor/ Model legend refers to the factor corresponding to the differentially expressed gene set and the DESeq2 model used (no pesticide-hatched bars; OP or Pyr- solid bars; BtTx = main effect of *Btt* treatment; OP PTx = main effect of OP exposure; OP or Pyr PTx:BtTx = interactions between OP or Pyr exposure and *Btt* treatment; Pyr Reg:BtTx: interaction between Pyr selection regime and *Btt* treatment). Colored circles on the left indicate transcripts included in the magenta WGCNA module (Suppl. Fig. 7). Transcript abbreviations are as follows: Stc = shuttle craft-like protein, Def = defensin, Att = attacin, PR = pathogenesis-related protein, HRG = histidine-rich glycoprotein, SP = serine protease, P450 = cytochrome P450, CEH = carboxylic ester hydrolase, GST = glutathione S-transferase, ABC = ABC transporter, Chit = chitinase protein, CP = cuticle-related protein, PM = peritrophic matrix related protein, Vit = vitellogenin, Ins = insulin-like peptide, LSP = larval serum protein, AchE = acetylcholinesterase, KLF = Krüppel-like factor, ApoIII = apolipophorin-III, Neur = neuropeptide.

Only two transcripts (an elastin and one unannotated transcript) were upregulated upon *Btt* infection in OP-selection regime individuals relative to susceptible ones (Fig. 3A, no pesticide model OP Reg:BtTx interaction), indicating that evolved resistance to OP minimally affects the baseline response to infection in the absence of pesticides. Fourteen transcripts were differentially expressed in Pyr-selected individuals relative to susceptible-regime individuals after *Btt* infection (Fig. 3A; no pesticide model Pyr Reg:BtTx interaction), including the downregulation of two defensins, a pathogenesis-related protein, and a histidine-rich glycoprotein (Fig. 4B, no pesticide model Pyr Reg:BtTx). Notably, these genes were differentially expressed in the opposite direction in Pyr regime-*Btt* infected larvae compared to gene expression induced by the main effect of *Btt* exposure, indicative of dampened expression of these genes (Fig. 4B, no pesticide model Pyr Reg:BtTx vs. BtTx, *e.g.* Def: TC006250). GO enrichment analysis revealed significant enrichment of humoral immune response and defense response among the differentially expressed genes from this interaction term supporting the impact on immune processes associated with the interaction effect of Pyr-selection regime with *Btt* infection (Suppl. Fig. 4A, Pyr Reg:BtTx).

### Effects of Pesticide Selection and Dual Exposure to Pesticides and Pathogens on Gene Expression

Next, we introduced the pesticide-exposed samples into the analysis, still treating OP and Pyr-associated treatments separately, to investigate interactions between pesticide selection regime, pesticide exposure, and *Btt* exposure on gene expression. For both pesticide class analyses, pesticide exposure had the greatest effect on gene expression (Figs. 3B-C; Pesticide Tx OP or Pyr) and induced differential expression of many canonical detoxification genes, including several cytochrome P450s, esterases, GSTs, UDP-glucuronosyltransferases (UGTs), and ABC transporters (Suppl. Fig. 5; OP or Pyr PTx). Genes differentially expressed after both OP and Pyr exposure were commonly enriched for GO processes involved in xenobiotic and detoxification responses including oxidoreductase and serine hydrolase activity (Suppl. Fig. 4B).

Genes differentially expressed with respect to pesticide selection regime and to the interaction between selection regime and pesticide treatment can identify genes associated with pesticide resistance. Overall, Pyr-regime had a greater effect on differential gene expression compared to OP-regime (Fig. 3; Regime OP or Pyr). Similarly, a greater number of genes were differentially expressed with the interaction between Pyr regime and Pyr exposure compared to OP regime and OP exposure (Figs. 3B [Regime OP, Pesticide Tx OP], 3C [Regime Pyr, Pesticide Tx Pyr]). Both OP-regime and Pyr-regime induced differential expression of canonical detoxification (e.g., P450s, esterases, GSTs, and ABC transporters) and cuticle-associated genes (Suppl. Figs. 5, 6; OP or Pyr Reg). GO enrichment analyses revealed the importance of structural constituent of cuticle for genes differentially expressed with OP regime and the interaction between OP regime and OP exposure (Suppl. Fig. 4B; OP Reg and OP Reg:PTx). Pyr-regime individuals differentially expressed genes enriched for molecular functions involved in oxidoreductase activity, heme binding, and structural constituent of cuticle, while genes differentially expressed with the interaction between Pyr regime and Pyr exposure were enriched for these functions and also serine hydrolase activity (Suppl. Fig. 4B; Pyr Reg and Pyr Reg:PTx). Thus, GO enrichment analyses revealed the common differential expression of cuticle-related genes in both OP and Pyr resistance, while Pyr resistance was additionally associated with differential expression of oxidoreductase metabolic detoxification processes.

In the models containing pesticide exposure samples, no genes were differentially expressed with the interaction between OP regime and *Btt* treatment (Fig. 3B; Regime OP, Bt Tx +) providing further evidence that, here, evolved OP resistance had minimal effects on immune responses to *Btt*. In contrast, 61 genes were differentially expressed with the interaction between Pyr selection regime and *Btt* treatment (Fig. 3C; Regime Pyr, Bt Tx +), although there was no significant GO term enrichment. Mirroring patterns observed in the no-pesticide model, the model that includes pesticide exposure samples also indicates that Pyr regime larvae had lower induced expression of a defensin-like transcript compared to control-regime larvae when exposed to *Btt* (Fig. 5B).

**Figure 5.**
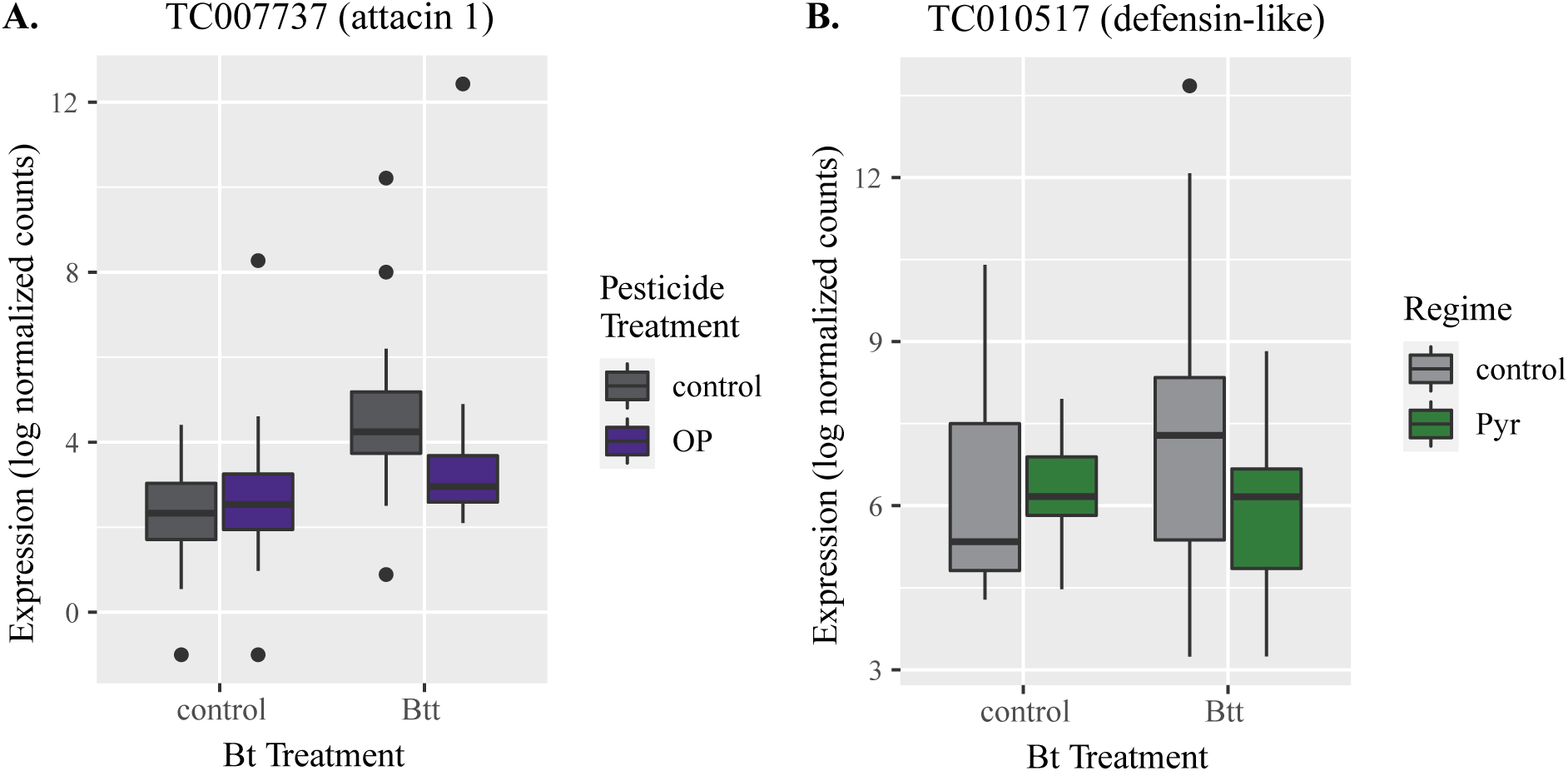
Normalized gene counts illustrate dampening effects of Pyr selection regime and OP treatment on immune gene expression induced with Bt expsoure. **A.** OP (OP model- OP PTx) and Bt treatment (OP model- BtTx) effects are shown for TC007737 (attacin1), and **B.** Pyr Regime (Pyr model- Pyr Reg) and Bt treatment (Pyr model- BtTx) effects are shown for TC010517 (defensin-like).

When considering the interaction effect between pesticide exposure and *Btt* infection, 175 genes were differentially expressed with dual OP and *Btt* exposure (Fig. 3B; Pesticide Tx OP, Bt Tx +), and only one gene (an upregulated pupal cuticle protein) was differentially expressed with dual exposure to Pyr and *Btt* (Fig. 3C; Pesticide Tx Pyr, Bt Tx +). OP pesticide exposure interacted with *Btt* exposure to induce differential gene expression of immune, detoxification, cuticle, and development genes compared to expression induced by the main effect of *Btt* exposure (Fig. 4A; compare BtTx and OP PTx:BtTx). Further analysis revealed that these interaction effects were mainly driven by significantly dampened upregulation upon combined OP and *Btt* exposure compared to upregulation induced with *Btt* alone; transcripts upregulated with *Btt* alone were not differentially expressed or were downregulated with combined OP and *Btt* exposure (Fig. 4A). For example, while *Btt* exposure alone induced upregulation of two attacins, three cytochrome P450s, and several cuticular proteins, larvae exposed to both OP and *Btt* had more modest differential expression of these same transcripts (e.g. *attacin-1* in Fig. 5A). Genes differentially expressed with the interaction between OP treatment and *Btt* exposure were enriched for GO terms involved in the molecular functions of oxidoreductase activity and structural constituent of cuticle (Suppl. Fig. 4B; OP PTx:BtTx).

### Correlated patterns of gene expression using WGCNA

Weighted gene correlation network analysis (WGCNA) allows for the identification of genes with expression correlated and significantly associated with experimental factors. We identified 25 modules containing genes with highly correlated expression, and several of these modules significantly correlated with one or more experimental factors (OP Regime, Pyr Regime, OP PTx, Pyr PTx, BtTx; Suppl. Fig. 7). We mapped differentially expressed transcripts to each gene module and identified one that was significantly associated with all five experimental factors (Suppl. Fig. 7; magenta module) and was also associated with a high number of significantly differentially expressed transcripts (Fig. 4; Suppl. Fig. 7; 201 DE transcripts/ 258 total). Thus, genes in this module displayed common expression patterns and were overall upregulated with both OP and Pyr selection regimes and downregulated with OP, Pyr, and *Btt* exposure (Suppl. Fig. 7). GO enrichment analysis of the magenta module transcripts revealed the importance of genes involved in hydrolase activity (GO:0016787; FDR = 6.07*10^-7^) and monooxygenase activity (GO:0004497; FDR = 1.81*10^-6^).

## Discussion

Our study experimentally evolved pesticide resistance in flour beetles (*T. castaneum*) to investigate the interactive effects of pesticide resistance and pesticide exposure on immunity and host-pathogen interactions. Pesticide-resistant insects and insects exposed to *Btt* both developed more slowly than their counterparts, indicating sub-lethal fitness costs to resistance and infection, respectively. We found that Pyr-resistant insects exhibited dampened transcriptional immune responses to *Btt* infection relative to their control-regime counterparts, although overall survival and development did not differ among OP- and Pyr-resistant and susceptible insects after *Btt* exposure. While the evolution of resistance did not substantially influence host infection phenotypes, exposure to pesticides did. Specifically, we observed that larvae were more likely to survive OP exposure if they were also orally exposed to *Btt*, with gene expression patterns suggesting dampened transcriptional stress responses in those beetles. Taken together, our data suggest that evolved resistance, pesticide exposure, and biopesticide infection all exert contrasting effects on life history parameters that could ultimately influence the demography and control of pest populations in complex ways. The factorial nature of our study design, moreover, provides a new window into the physiological basis and fitness consequences of host responses to overlapping environmental stressors.

Insects, including *T. castaneum*, can rapidly evolve resistance to pesticides through mechanisms that enhance metabolic detoxification and cuticular modifications (Rösner et al., 2020). Here, populations selected for pesticide resistance evolved it within 6-8 generations (Birnbaum et al., 2021), demonstrating increased survival and faster development after pesticide exposure compared to control-selected populations. Differential gene expression between pesticide- and control-selection regimes suggested that OP resistance is associated with constitutive changes in cuticle gene expression while Pyr resistance is associated with oxidoreductase activity, heme binding, and cuticular modifications, consistent with previous studies in OP and Pyr resistant insects (Carvalho et al., 2013; Tandonnet et al., 2020; Zimmer et al., 2017). Thus, while both OP and Pyr resistance involved changes in cuticle gene expression, Pyr resistance was also associated with changes in metabolic detoxification.

Oral *Btt* infection did not decrease survival in our study but was associated with slower development in *T. castaneum* larvae, indicating a direct cost of infection or an indirect cost from mounting an immune response, as previously observed in the same system (Tate & Graham, 2015) and in other insects challenged with Bt toxins (Hernández-Martínez et al., 2017; Rahman et al., 2011). Previously documented variation in resistance in *T. castaneum* against Cry3 proteins effective against other Tenebrionid beetles may explain why *Btt* had negative effects on development but did not instigate high mortality (Milutinović et al., 2013; Oppert et al., 2011). Here, exposure to *Btt* induced the upregulation of AMPs (attacins and defensins) and one pathogenesis-related protein as well as differential expression of detoxification genes, including P450s and a CEH, GST, an ABC transporter, and several cuticle and development related genes. Previous transcriptomic studies of oral Bt infection in *T. castaneum* have identified similar transcripts, including attacin and defensin AMPs, a pathogenesis-related protein, a P450, and a GST (Behrens et al., 2014; Greenwood et al., 2017), supporting the assumption that *Btt* oral exposure in our experiments presented an immune challenge despite the lack of infection-induced mortality.

Previous work has suggested that exposure to OP and Pyr pesticides can reduce defenses against pathogens and stimulate an array of changes in humoral, cellular, and oxidative stress components of immunity (Dubovskiy et al., 2014; Gu et al., 2016; Hauser & Koella, 2020). Our results reveal a strong interaction effect between *Btt* and OP exposure that manifests at both phenotype and gene expression levels. For example, dual oral exposure to *Btt* and OP improved larval survival and partially rescued development time relative to OP exposure alone, without exacerbating *Btt*-induced mortality. At the same time, *Btt* co-exposure modified the differential expression of 175 genes relative to OP exposure alone (Fig. 3B). In contrast, Pyr exposure did not influence survival with *Btt* infection, but larvae exposed to both Pyr and *Btt* also partially rescued development time compared to those exposed only to pesticides. However, in contrast to the effects on gene expression observed with dual OP and *Btt* exposure, *Btt* co-exposure with Pyr had negligible effects on gene expression with the overexpression of only one cuticle-related transcript. It is possible that the positive effects on development time seen with pesticide and *Btt* co-exposure compared to single exposure to pesticides are part of an advantageous response to increased risk (Roth & Kurtz, 2008) as has been seen in viral infected caterpillars reared on more toxic diets (Smilanich et al., 2018).

There is also the potential for indirect antagonistic interactions between OP and *Btt*, e.g. through induction of cross-protective host defenses. For example, while *Btt* exposure induced upregulation of attacins, cytochrome P450s, and cuticle-related transcripts, combined OP and *Btt* exposure dampened the expression of these transcripts. It is possible that this dampened expression of immune and detoxification genes may underlie reductions in immunopathology or ROS damage corresponding to increased fitness, or the benefits to fitness may be due to differences in resource allocation when faced with dual pressures (Lazzaro & Tate, 2022). Interestingly, single exposure to OP or *Btt* induced downregulation of an ABC transporter but this transcript was upregulated with co-exposure to OP and *Btt*. ABC transporters have been identified as receptors for Bt Cry proteins and aid in membrane pore formation (Güney et al., 2021; Heckel, 2021; Pauchet et al., 2016), and the additional function of ABC transporters as xenobiotic pumps may help to mediate Bt toxicity (Dermauw & Leeuwen, 2014; Wu et al., 2019). It is also possible that OP and *Btt* interact directly, as would be the case if *Btt* actively degrades OP for energy (Kumar et al., 2018) or if OP facilitates *Btt* germination rates. Our *in vitro* data suggest that *Btt* vegetative cells in liquid medium replicate faster in the presence of OP, lending credence to a direct interaction. On the other hand, there was no difference in germinated spore counts between control-diet and OP-diet disks, suggesting that direct interactions, if they matter *in vivo*, are not relevant until after the host ingests the diet.

Supporting the idea of localized cross-protective mechanisms, a previous study of oral vs. septic *Btt* infection in *T. castaneum* demonstrated that oral infection induced greater changes in cuticle-related gene expression compared to septic infection (Behrens et al., 2014). In our study, OP-exposed individuals were not protected from greater mortality upon septic *Btt* infection, suggesting that infection route has important implications for pesticide-pathogen interactions. We found cuticle-associated gene expression changes induced by OP exposure, and cuticular modifications of gut tissues may provide an additional layer of protection from oral Bt infection (Raymond et al., 2010). Future work dissecting the relative importance of these direct and indirect interactions will be crucial for refining integrative pest management (IPM) strategies employing both chemical and biopesticides.

The impact of evolved pesticide resistance on immunity is still poorly understood (Minetti et al., 2020), and in the absence of pesticides we did not observe any significant interactions between pesticide resistance and Bt exposure on survival or development. Few genes were differentially expressed between OP resistant and susceptible beetles after *Btt* exposure, but *Btt* exposure in Pyr-resistant populations was associated with dampened upregulation of several genes, including two defensins, a histidine-rich glycoprotein, and a pathogenesis-related protein, that were more strongly upregulated in Pyr-susceptible populations after *Btt* exposure. It is also possible that Pyr-resistant populations had constitutively higher immune gene expression resulting in reduced immune activation upon Bt infection. Moreover, an apolipophorin-III transcript, previously shown to be important in immunity against coleopteran-specific Bt toxins (Contreras et al., 2013), was upregulated in Pyr-resistant populations. In contrast, cuticle-related protein and neuropeptide transcripts were downregulated in Pyr-regime larvae exposed to *Btt*. These immune gene changes associated with Pyr resistance may have important effects on interactions with other pathogens, and future research into Pyr resistance interactions with other pathogens may yield important insights.

While we did not observe any phenotypic differences with pesticide selection regime and *Btt* infection here, we also did not observe mortality from *Btt* alone and it is possible that the immune challenge was not strong enough to elicit an interaction with resistance mechanisms. However, the differential expression of immune and cuticle functions in Pyr-regime larvae exposed to *Btt* could have important implications for insect interactions with other pathogens or xenobiotic pressures by modifying detoxification capabilities or providing an additional barrier against infection or toxins (Erlandson et al., 2019; James & Xu, 2012; Lilly et al., 2016). Furthermore, changes in oxidoreductase activity observed after Pyr selection are likely to alter the status of reactive oxygen species (ROS) within the host and may impact immunity against different pathogens (Minetti et al., 2020; Wang et al., 2016). Finally, it is possible that adaptation to pesticides could constrain host evolutionary responses to pathogens, or vice versa. Future experiments could simultaneously select for dual resistance to both pesticides and pathogens to see if evolutionary trajectories are similar or constrained relative to selection under a single stressor.

### Conclusions

A better understanding of the complex interactions between pesticide exposure and resistance for host-pathogen outcomes is critical for effective pest and vector control strategies (James & Xu, 2012; Minetti et al., 2020; Orr et al., 2020). Here, we identified antagonistic interactions between Bt and OP that mitigated the effects of pesticides on fitness-related host traits. Our results suggest that both pesticides and Bt individually influence important demographic parameters like development time and mortality rate, but that co-exposed hosts may actually perform better than those exposed to pesticides alone. We also observed dampened expression of important candidate genes upon dual exposure to OP and *Btt* and in Pyr-resistant individuals exposed to *Btt* compared to the main effect of *Btt* exposure. Are these within-host interaction effects sufficient to alter the dynamics of disease and economic damage in pest populations? Mathematical models would be useful to address this question, as the answer will likely depend on feedbacks from ecological variables like intraspecific competition, relative transmission among single- and co-exposed hosts, and the relative costs of pesticides and infection to host reproductive rates (Booton et al., 2019; Booton et al., 2018; Kuniyoshi & Santos, 2017). Moreover, our data suggest that the rise of pesticide resistance within populations could reduce the magnitude of the interaction effects, likely altering population outcomes. Overall, our results shine light on the physiological and evolutionary processes that shape interactions among two important ecological stressors and should prompt further study of their interactions at multiple levels of biological organization on ecological and evolutionary time scales.

## Data Accessibility

Sequence reads are deposited in the NCBI SRA under the BioProject Accessions: PRJNA753142, PRJNA756317.

## Acknowledgements

We would like to thank three anonymous reviewers for comments that helped improve the manuscript. We would like to thank William Galardi for assistance in preliminary dose response experiments and Siqin Liu for assistance with population maintenance. We are also grateful to Justin Critchlow and Justin Buchannan for guidance with *in vitro* bacterial growth curves. Last, we would like to thank Juan Luis Jurat-Fuentes for comments on the manuscript. This work is funded by USDA NIFA 2019-67012-29659 provided to S.S.L.B. N.K.E.S. and A.T.T. were supported in part by Alfred P. Sloan Foundation Fellowship FG-2020-12949 to A.T.T.

## Supplemental Table Legends

**Suppl. Table 1.** Read counts and alignment statistics for RNAseq libraries.

**Suppl. Table 2.** Significantly differentially expressed (DE) transcripts from the no pesticide model (Conly), OP model (allOP), and Pyr model (allPyr) (p < 0.05). Fold change differences (log2FC) and adjusted p-values (padj) are listed for each factor within each model (BtTx: main effect of Bt; OPReg, PyrReg: main effect of OP and Pyr selection regime, respectively; OPPTx, PyrPTx: main effect of OP and Pyr pesticide treatment; OPReg:PTx, PyrReg:PTx: interaction effect between pesticide selection regime and exposure; OPReg:BtTx, PyrReg:BtTx: interaction effect between selection regime and Bt expsoure; OPPTx:BtTx, PyrPTx:BtTx: interaction effect between pesticide and Bt exposure. DE transcripts belonging to WGCNA modules are listed (moduleColor).

**Suppl. Table 3.** Main and interaction effects of pesticide exposure, selection regime, and Bt exposure on survival and adult development (coxme(Surv(Day, Status) ∼ PTx+ Regime + BtTx + Reg:PTx + Reg:BtTx + PTx:BtTx +(1|Pop)); compared to control regime and treatments).

**Suppl. Table 4.** Effect of pesticide exposure on in vitro Bt growth. The effects of OP and Pyr treatment were analyzed separately. The effect of treatment was analyzed for each; if replicate was non-significant, all replicates were combined.

**Suppl. Table 5.** Effect of pesticide treatment on germinated spores from oral diet disks.

**Suppl. Table 6.** Survival after septic Bt infection.

## Supplemental Figure Legends

**Suppl. Figure 1.**
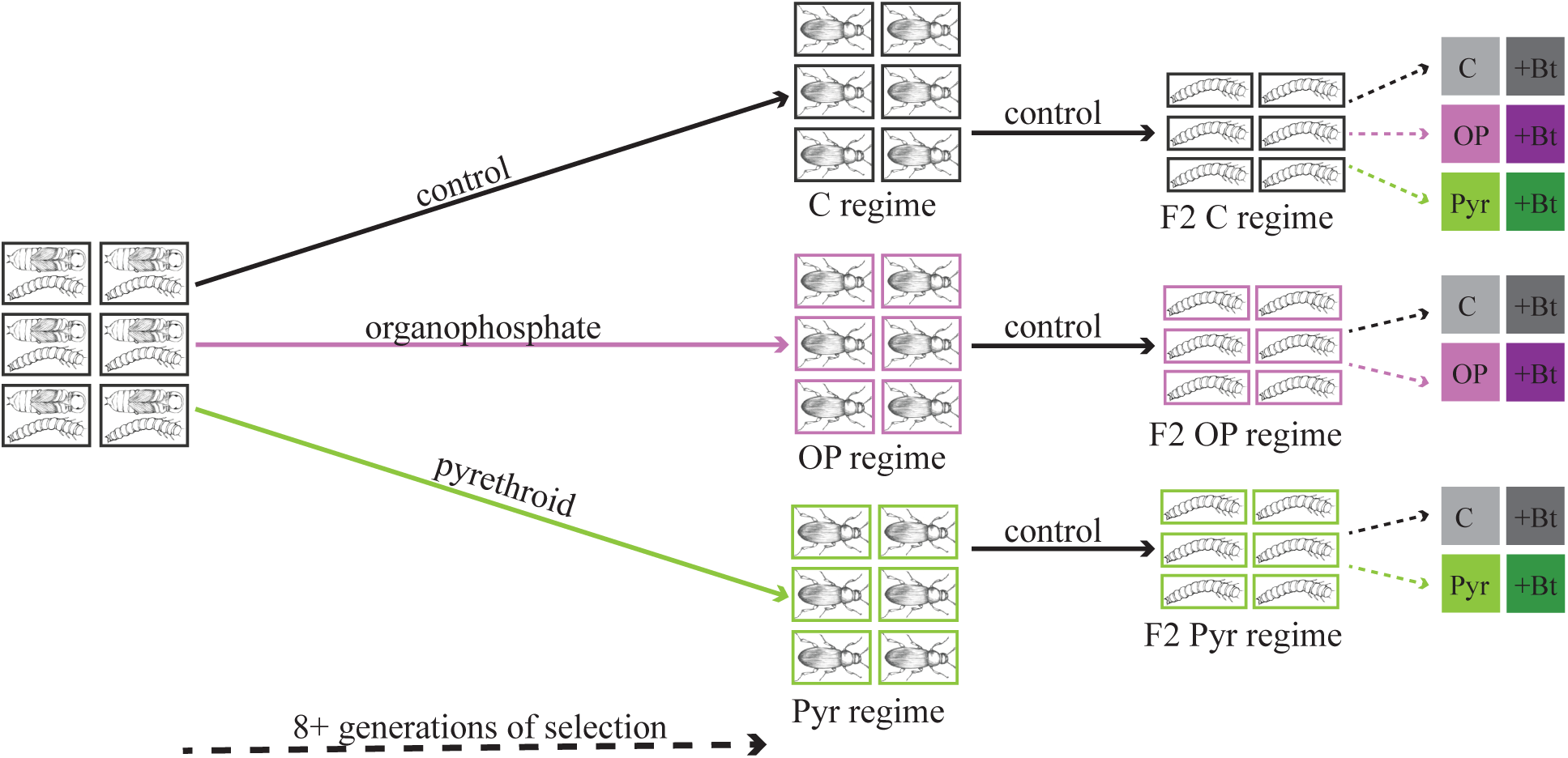
Experimental design schematic. Larvae from six distinct populations were exposed to control, organophosphate (OP), or pyrethroid (Pyr) diets for at least eight generations. Adults from Gen. 8 pesticide-selected and control populations were allowed to reproduce on control diets, and twelve-day old F2 larvae from each population were exposed to control and pesticide diets with and without *Btt*.

**Suppl. Figure 2.**
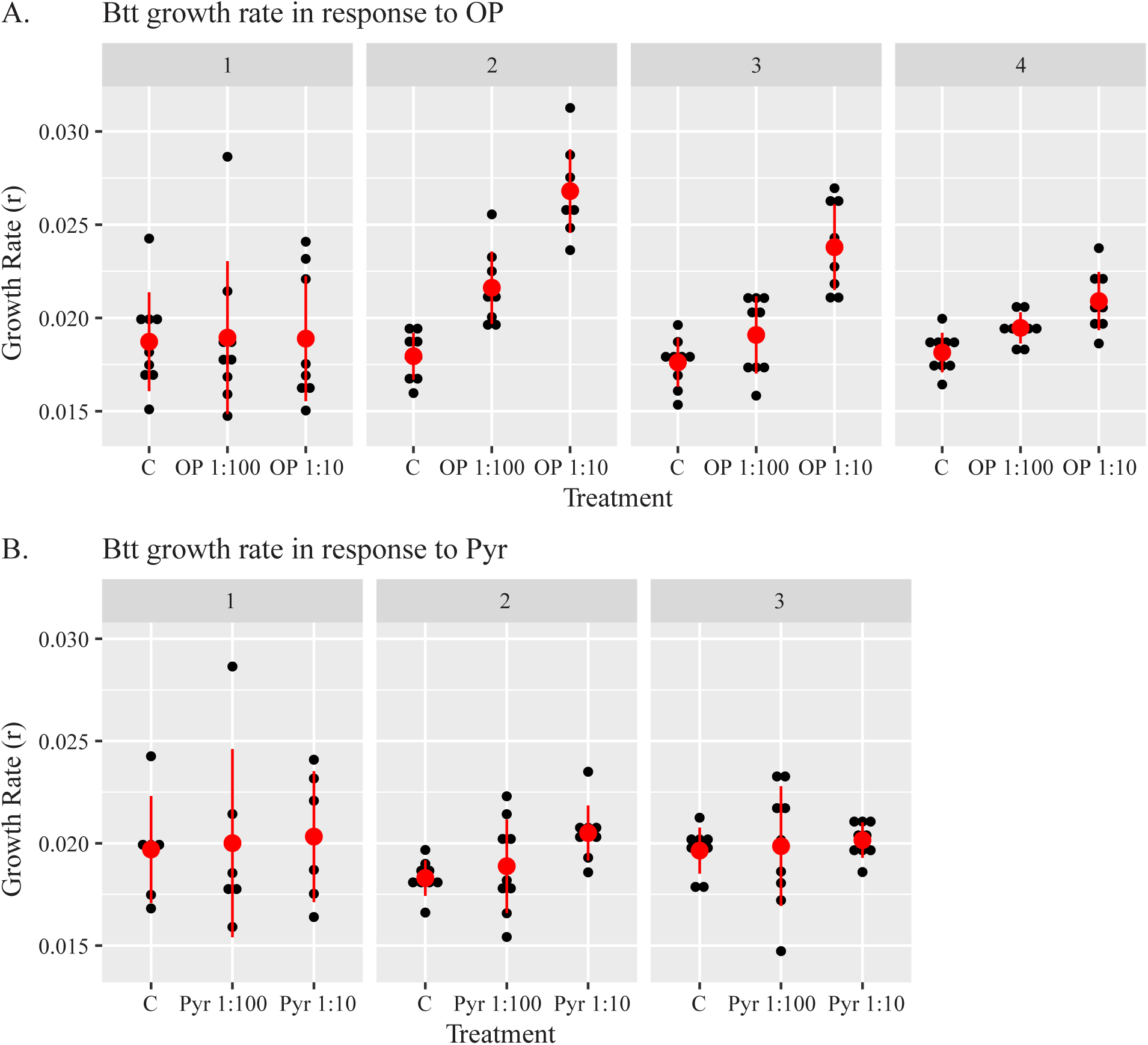
*In* vitro growth rate (r) of *Btt* under control and pesticide conditions. **A.** *In vitro* growth rate (r) of *Btt* under control and OP conditions. Data were not normally distributed, so non-parametric tests were used. There was a significant difference between replicates, so the effect of treatment was analyzed for each replicate separately (Suppl. Table 4). K-W test revealed significant positive effect of OP on *Btt* growth rate for three of the four replicates (reps 2-4; Suppl. Table 4). **B.** *In vitro* growth rate (r) of *Btt* under control and Pyr conditions. Data were not normally distributed, so non-parametric tests were used. There was no significant difference between replicates or with the effect of treatment (Suppl. Table 4).

**Suppl. Fig. 3.**
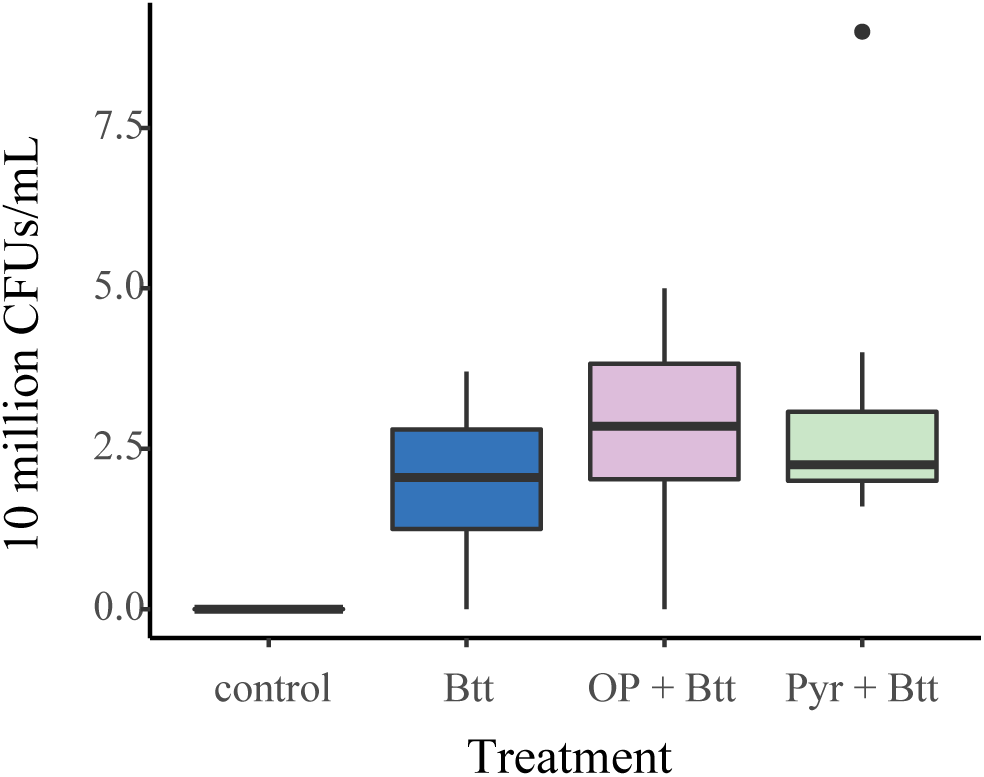
Germination of spores from diet disks in the absence or presence of either pesticide (Bt, OP + *Btt* or Pyr + *Btt*: n=10, control: n=6). There was no significant difference between Bt control and Bt pesticide diet disks (Suppl. Table 5).

**Suppl. Figure 4.**
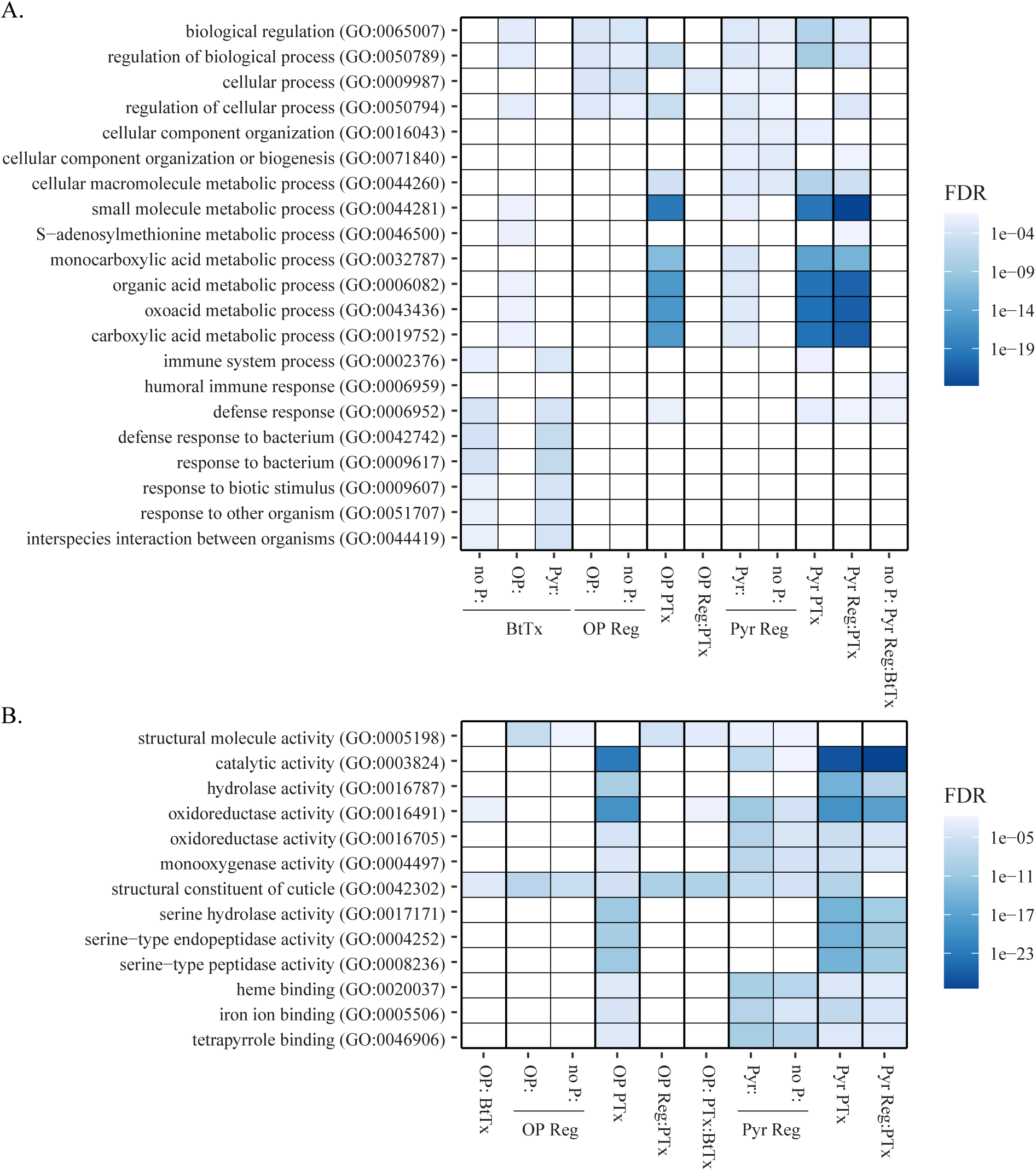
Top significantly enriched GO terms for differentially expressed gene sets. **A.** Biological process GO terms, **B.** Molecular function GO terms. Differentially expressed gene sets are denoted on the x-axis and refer to the DESeq2 model (no P: = no pesticide model, OP: = OP model, Pyr: = Pyr model) and the factor corresponding to the differentially expressed gene set (BtTx = main effect of *Btt* treatment; OP Reg, Pyr Reg = main effect of OP and Pyr selection regimes, respectively; OP PTx, Pyr PTx = main effect of OP and Pyr exposure, respectively; OPReg:PTx, PyrReg:PTx = interaction between OP and Pyr regime and exposure, respectively; PyrReg:BtTx = interaction between Pyr regime and *Btt* treatment; OPTx:BtTx = interaction between OP exposure and *Btt* treatment).

**Suppl. Fig. 5.**
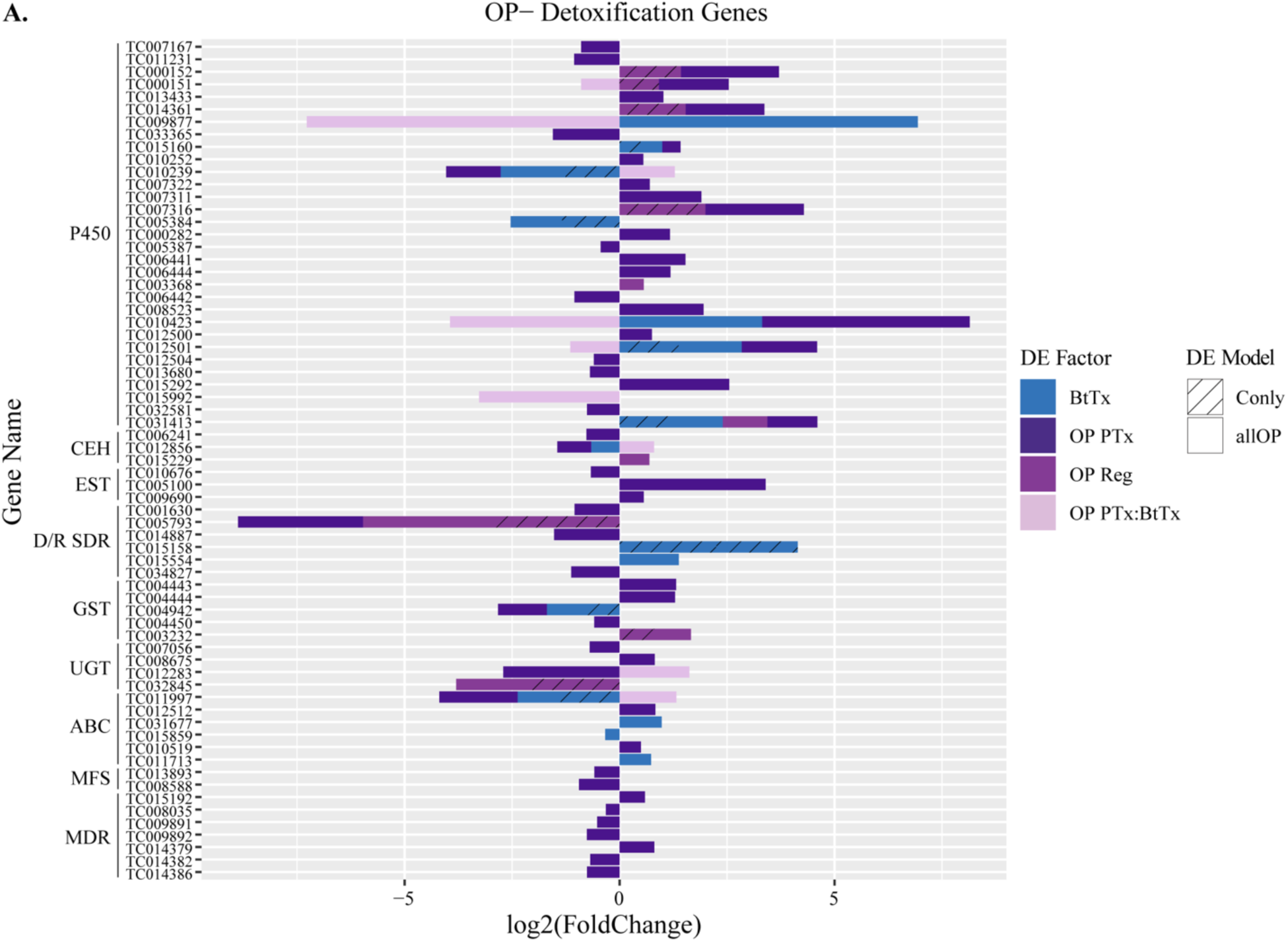
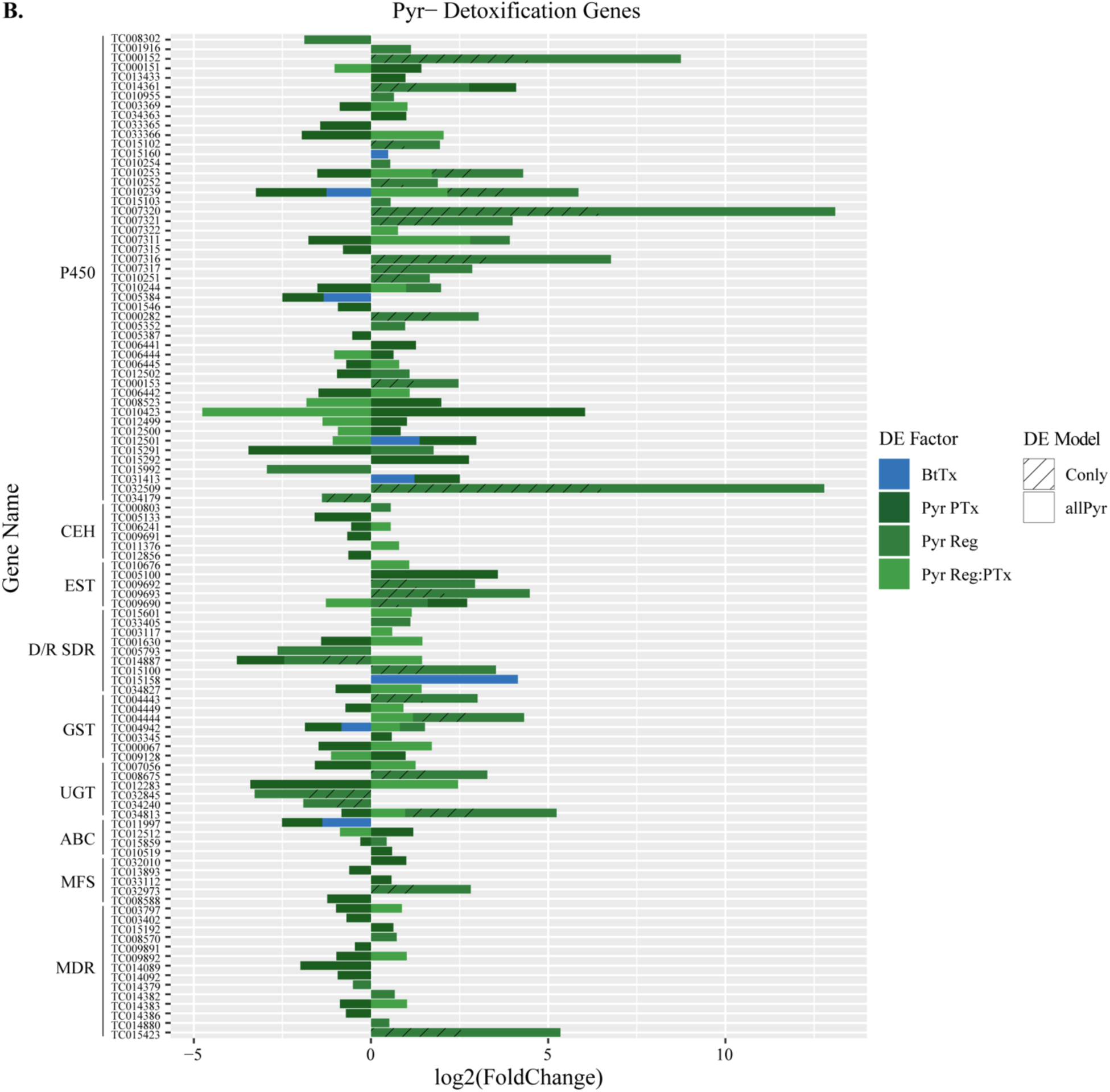
Canonical detoxification genes significantly differentially expressed (padj < 0.05) with **A.** OP exposure (OP PTx), OP selection regime (OP Reg), *Btt* exposure (BtTx), and the interaction between OP regime and exposure (OP Reg:PTx) and **B.** Pyr exposure (Pyr PTx), Pyr selection regime (Pyr Reg), *Btt* exposure (BtTx), and the interaction between Pyr regime and exposure (Pyr Reg:PTx).

**Suppl. Fig. 6.**
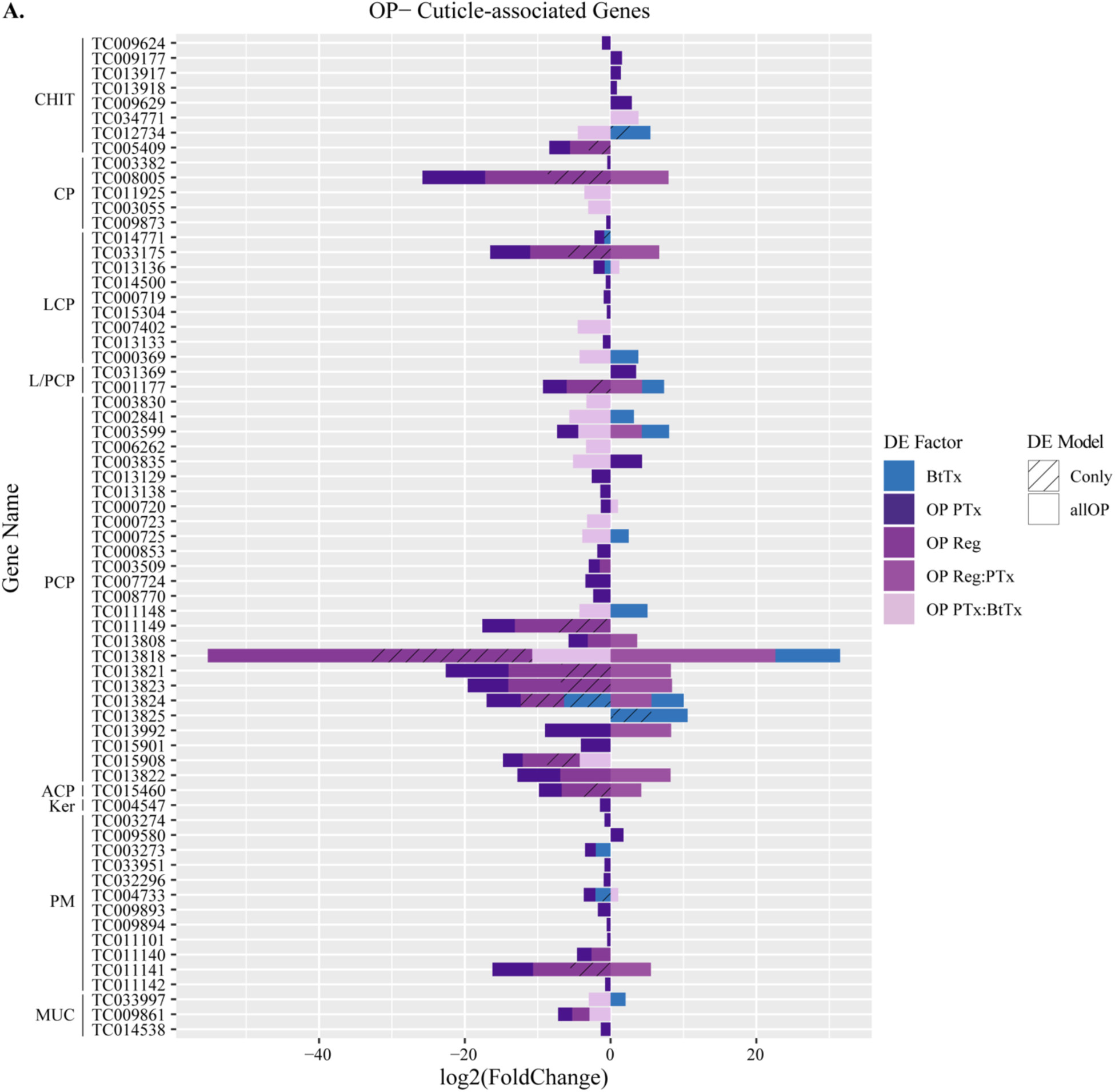
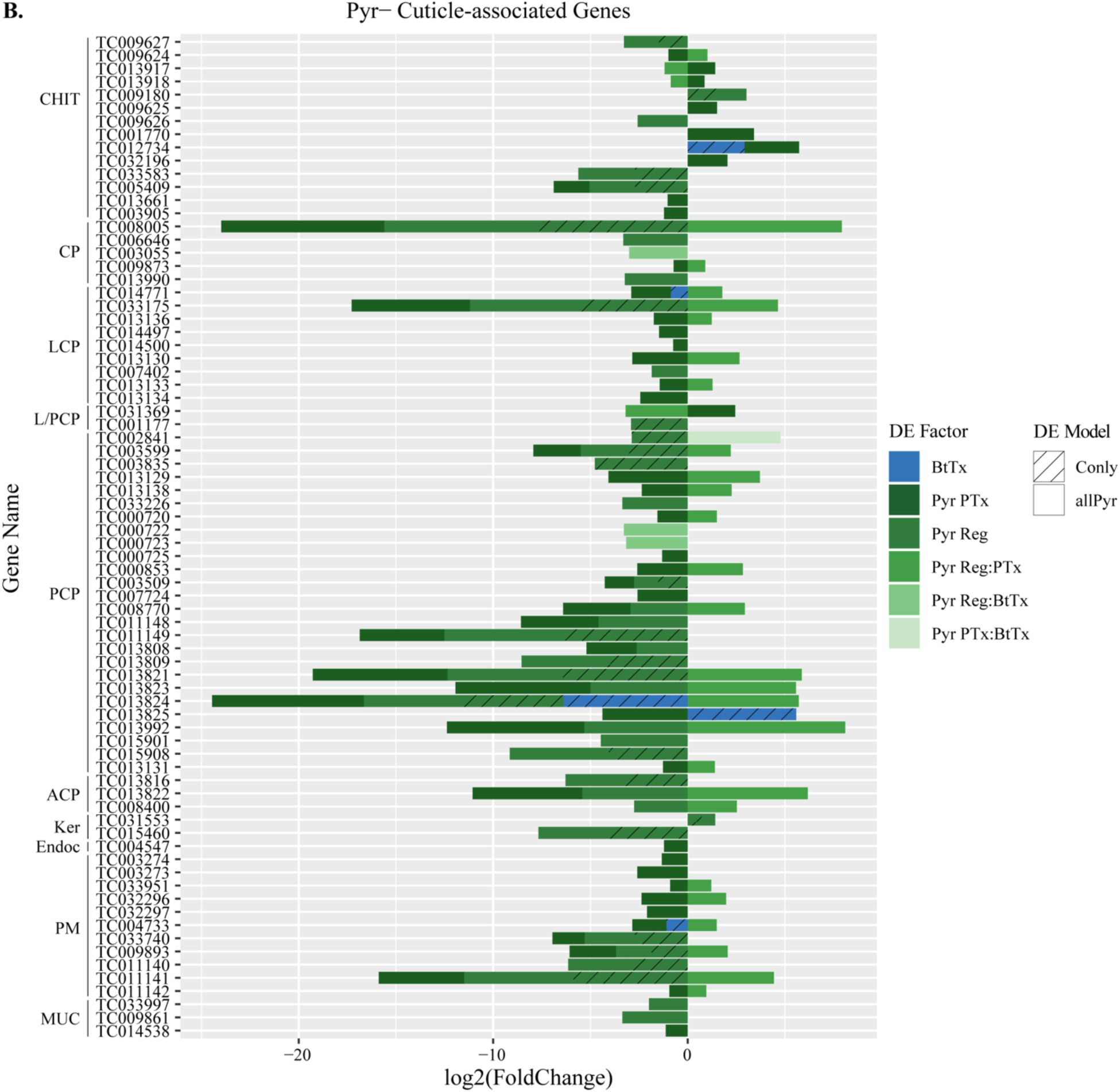
Cuticle-associated genes significantly differentially expressed (padj < 0.05) with **A.** OP exposure (OP PTx), OP selection regime (OP Reg), *Btt* exposure (BtTx), the interaction between OP regime and exposure (OP Reg:PTx), and the interaction between OP and *Btt* exposure (OP PTx:BtTx), and **B.** Pyr exposure (Pyr PTx), Pyr selection regime (Pyr Reg), *Btt* exposure (BtTx), the interaction between Pyr regime and exposure (Pyr Reg:PTx), the interaction between Pyr regime and *Btt* exposure (Pyr Reg:BtTx), and the interaction between Pyr and *Btt* exposure (Pyr PTx:BtTx).

**Suppl. Fig. 7.**
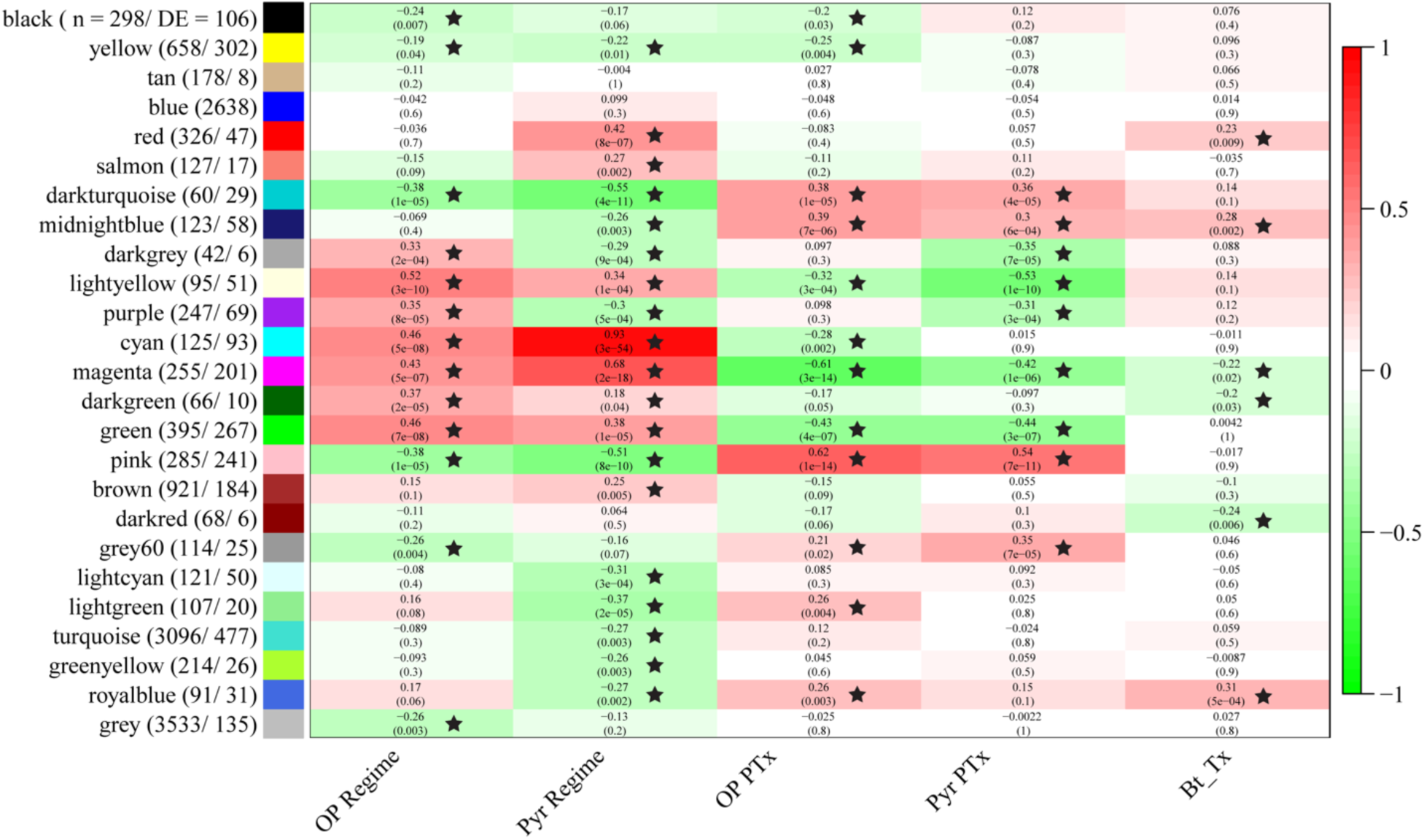
WGCNA gene modules and significant relationships with experimental factors. The total number of transcripts (n) and number of differentially expressed (DE) transcripts in each module are listed, and significant relationships with experimental factors are indicated with black stars. Top values within each cell indicate the correlation strengths (corresponding to the heat map colors) and bottom values in parentheses indicate p-value significance for the relationship of each module with the different experimental factors. Significant (p < 0.05) positive correlation values and red cells indicate that the transcripts within the module are commonly upregulated; negative significant correlation values and green cells indicate that the transcripts within the module are commonly downregulated.

## Notes

### Competing Interest Statement

The authors have declared no competing interest.

